# Myths and mechanisms: RecBCD and Chi hotspots as determinants of self *vs*. non-self

**DOI:** 10.1101/2021.07.08.451572

**Authors:** Suriyen Subramaniam, Gerald R. Smith

## Abstract

Bacteria face a challenge when DNA enters their cells by transformation, mating, or phage infection. Should they treat this DNA as an invasive foreigner and destroy it, or consider it one of their own and potentially benefit from incorporating new genes or alleles to gain useful functions? It is frequently stated that the short nucleotide sequence Chi (5’ GCTGGTGG 3’) recognized by RecBCD helicase-nuclease allows *Escherichia coli* to distinguish self (*i.e., E. coli*) DNA from non-self (*i.e.*, any other) DNA and to destroy non-self DNA, and that Chi is “over-represented” in the *E. coli* genome. We show here that these dogmas are incorrect and apparently based on false assumptions. We note Chi’s wide-spread occurrence and activity in distantly related species. We illustrate multiple, highly non-random features of the genomes of *coli* and coliphage P1 that account for Chi’s high frequency and genomic position, leading us to propose that P1 selects for Chi’s enhancement of recombination, whereas *E. coli* selects for the preferred codons in Chi. We discuss other, substantiated mechanisms for self *vs.* non-self determination involving RecBCD and for RecBCD’s destruction of DNA that cannot recombine, whether foreign or domestic.

## Introduction

Potentially dangerous DNA frequently enters bacterial cells *via* various mechanisms. During transformation, naked DNA enters cells, in some species by specialized, high-efficiency up-take mechanisms, and can be integrated into the recipient genome by recombination. Phages actively inject their DNA into cells to initiate infection, which can kill the cells. Rarely, the phage particle contains bacterial DNA incorporated from the previous host’s genome; the injected DNA, as in transformation, can either integrate into the recipient’s genome or set in motion destructive acts. Conjugative plasmids encode specialized machinery to transfer DNA directly into the recipient cell, thereby bypassing exposure of the DNA to a potentially destructive environment. In some cases, the plasmid also transfers bacterial chromosomal DNA along with the plasmid DNA.

In each case, the imported DNA can come from a bacterial species different from the recipient and can be considered “foreign.” DNA transfer can be directly observed in the laboratory, and these transfers likely occur in nature, too. It is inferred to occur frequently in nature from the presence, in unrelated species, of DNA “islands” – highly conserved DNA sequences with one or more genes, often functionally related indicating they travel together as a unit (Hacker and Kaper, 2000). Foreign, or non-self, DNA is more likely to be deleterious than self DNA, and it is reasonable that bacteria have ways to distinguish the two. They can profit from some DNA (self as well as beneficial non-self DNA such as islands encoding drug-resistance), but they are at risk of imminent death from other DNA (non-self).

How, then, do bacteria, such as *E. coli,* distinguish self from non-self DNA? Numerous authors have stated that the short DNA sequence called Chi occurs at higher than expected frequency in the *E. coli* genome and is a signal that allows *E. coli* to tell self from non-self DNA [*e.g.*, (Biswas et al., 1995; Kuzminov, 1995; Anderson and Kowalczykowski, 1998; Kobayashi, 1998; El Karoui et al., 1999; Stahl, 2005; Levy et al., 2015)]. Chi (5’ GCTGGTGG 3’) is recognized by the *E. coli* RecBCD helicase-nuclease as it unwinds DNA rapidly [∼1000 base-pairs (1 kbp)/sec] and prompts RecBCD to nick the DNA at Chi and promote recombination between that DNA and other homologous double-stranded (ds) DNA in the cell (Smith, 2012) (see Figure 1 and discussion below for further explanation). RecBCD’s activity can spur incorporation of incoming self DNA and aid evolution; it can also repair broken DNA in a cell with multiple copies of its chromosome (as typical in growing bacteria). But in other situations, RecBCD’s nuclease activity can degrade DNA (Clark et al., 1966; Simmon and Lederberg, 1972). It is often stated that Chi instructs RecBCD to recombine self DNA (containing Chi) and to destroy non-self DNA (without Chi) [*e.g.*, (Touzain et al., 2011; Levy et al., 2015; Cheng et al., 2020)]. Below, we examine and show the weaknesses of three dogmatic views:

1. In *E. coli*, Chi distinguishes self from non-self DNA.
2. Chi occurs more frequently in *E. coli* DNA than expected.
3. Chi converts RecBCD from a destructive nuclease into a DNA break-repair machine.

**Figure 1.**
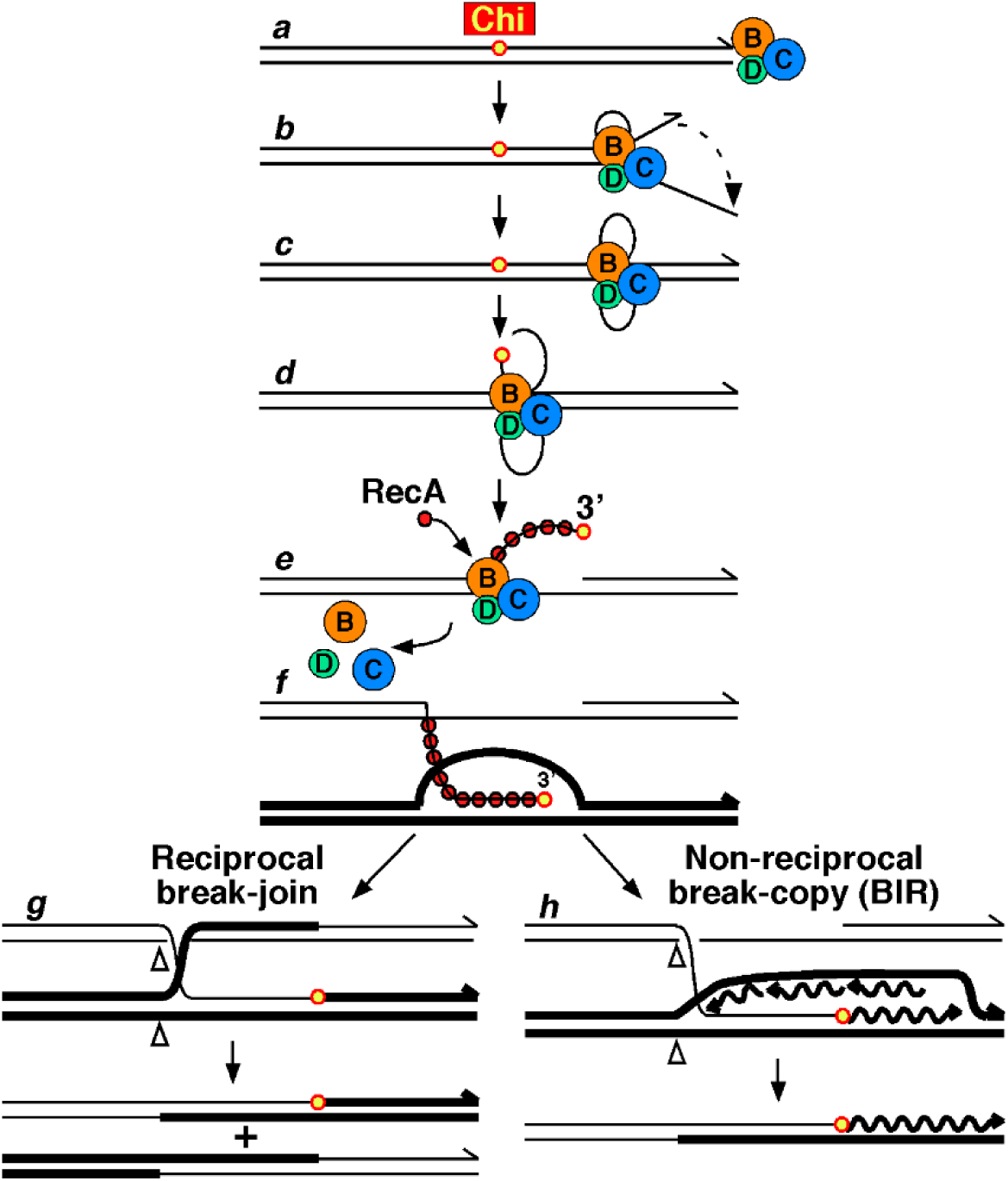
Pathway of recombination promoted by Chi and RecBCD helicase-nuclease. (***a***) RecBCD binds the end of ds DNA (from linear DNA entering the cell or from a chromosomal break). (***b***) The RecD helicase moves along the 5’-ended strand faster than the RecB helicase moves along the 3’-ended strand, forming a loop-tail structure. (***c***) The tails anneal to form a twin-loop (aka “rabbit ear”) structure. (***d***) Upon encountering Chi, the RecB nuclease cuts the 3’ ended strand about 5 nucleotides 3’ of the Chi octamer. (***e***) RecBCD loads RecA strand exchange protein onto the newly generated 3’-ended strand. (***f***) The RecA-ssDNA filament pairs with homologous dsDNA (thick line), such as the circular chromosome (only partially shown), forming a displacement (D-) loop. (***g***) The D-loop may be cut and form a Holliday junction, which can be resolved into reciprocal recombinants (cutting, swapping of ends, and ligation at open triangles). (***h***) Alternatively, the 3’ end in the D-loop may prime DNA synthesis and formation of a dual-strand replication fork, which can be resolved, as in ***g***, into one non-reciprocal recombinant (shown) and a parental-type (non-recombinant) DNA molecule (not shown). On linear DNA entering the cell, RecBCD acts as shown at both ends. Option ***g*** would generate a recombinant circle and a linear form, which could be lost or recombine again. Option ***h*** would generate two circles or a dimeric circle, which would be resolved by XerCD at the *dif* site into two circles (Smith, 1991). From (Amundsen et al., 2007) as modified from (Smith et al., 1981).

We then discuss Chi and RecBCD’s established physiological roles and mechanisms by which bacteria do discriminate between self and non-self DNA using RecBCD.

### Chi is more frequent in many bacterial species and *E. coli* phages and plasmids than in *E. coli* itself

The dogma that Chi is the signal for *E. coli* to distinguish its own DNA from that of other species or phages implicitly assumes that Chi occurs more frequently in *E. coli* DNA than in other DNA [number of Chi sites per million base-pairs (Mbp) of DNA; Chi’s “density”]. But this is not the case – the density of Chi is greater in many other bacterial species and *E. coli* phages than in *E. coli* itself (Table 1). For example, *E. coli* has 1008 Chi sites in its 4.64 Mbp genome (217/Mbp), but there are >300 Chi sites/Mbp in *Klebsiella pneumoniae* (another enteric bacterium, like *E. coli;* order Enterobacteriales) and in more distant *Pseudomonas aeruginosa* (order Pseudomonadales) and a *Xanthomonas* sp. (order Xanthomonadales). The dogma noted above would lead to the conclusion that *P. aeruginosa* is more like *E. coli* than *E. coli* itself. Even *Mycobacterium tuberculosis* (order Actinomycetales), very distant genetically and physiologically from the enteric bacteria, has higher Chi density (241/Mbp) than *E. coli*.

**Table 1.** Chi is frequent in foreign (“non-self”) DNA.

| Species | Chi sites/Mb | Rank of Chi among ~65,000* octamers |
| --- | --- | --- |
| *E. coli* phage P1 (temperate) | 527 | 1 |
| *Klebsiella pneumoniae* | 345 | 63 |
| *Pseudomonas aeruginosa* | 332 | 352 |
| *Xanthomonas* sp. | 330 | 350 |
| *E. coli* phage T3 (virulent) | 314 | 9 |
| *Serratia* sp. | 257 | 148 |
| *Mycobacterium tuberculosis* | 241 | 493 |
| **<u>Escherichia coli</u>** | **<u>217</u>** | **<u>21</u>** |
| *Salmonella enterica* | 179 | 145 |
| *Vibrio harveyi* | 110 | 290 |

\* Phages P1 and T3 have 55659 and 24755 unique octamers, respectively.
Bacteria listed have 65000 ± <1000 unique octamers each.
Data are from DNA sequences whose sources are in Table S9.

Some *E. coli* phages lack Chi, as the dogma would expect, but others have Chi, in some cases at two or three times higher density across their genome than in *E. coli* (Tables 1 and S1). This dogma likely derives, at least in part, from phage lambda, in which Chi was discovered, lacking Chi in its 48.5 kbp of DNA. But this had to be the case, for Chi was first observed as a mutation that increased the plaque-size of lambda deletion mutants lacking its Red recombination pathway and Gam inhibitor of RecBCD (Henderson and Weil, 1975). If lambda had Chi, Chi would not have been discovered in this way. Chi stimulates the *E. coli* RecBCD pathway to recombine monomeric circles of replicating lambda to form dimeric (or higher order concatemeric) DNA essential for packaging lambda DNA into viable phage particles (Lam et al., 1974; Stahl and Stahl, 1977). Without Chi, this recombination occurs, but at low level; consequently, lambda *red gam* mutant plaques are small. With Chi’s stimulation of recombination and phage propagation, the plaques are large. Note that the name Chi comes from its activity as a “crossover hotspot instigator” (Lam et al., 1974), but this key criterion is often overlooked when discussing Chi sites, as we note below. Study of Chi in lambda for 50 years has been essential to working out the complex activities of RecBCD and its control by Chi (Smith, 2012).

Remarkably, some phages closely related to lambda do have Chi (Bobay et al., 2013) (Table S1). For example, four phages with lambda-size genomes and lambda-like gene organization have Chi densities similar to or even greater than that of *E. coli*. Two of these four phages have apparent RecBCD inhibitors, like lambda’s Gam protein, and two do not. Most of these Chi sites are in the orientation of active Chi in lambda, implying they are active in these native phages (Chi is active as a recombination hotspot in lambda only when correctly oriented with respect to the direction of its DNA packaging; see Figure 1 and below). Other phages unrelated to lambda, including both temperate and virulent types, also have Chi. Especially notable is temperate phage P1 and three relatives (P7, D6, and putative phage NGUA22), which have Chi density two or three times higher than that of *E. coli.* These phages may well use Chi to their advantage (see section on temperate phage P1 below), but after most infections they kill *E. coli* despite having Chi. Three unrelated virulent phages (T3, T5, and T7) have Chi density near to or greater than that of *E. coli*. All these phages, obviously, succeed in killing *E. coli*: thus, Chi does not act as a signal of “self” DNA and thereby protect the cells from death. Even the virulent phages T2, T4, and T6 have Chi, yet they kill *E. coli* soon after infection. We see no discrimination by *E. coli* between phages with Chi and those without.

Many plasmids, which replicate independently of the chromosome, do not normally kill the cell when they enter by conjugation (Johnson and Grossman, 2015). Many, such as *E. coli*’s 99.2 kbp F (fertility) factor, lack Chi, yet they are beneficial to *E. coli* by providing a means for gene exchange and rapid evolution. But other conjugative plasmids have Chi at high density. For example, plasmid pMG252, which confers quinolone-resistance and was first isolated from *pneumoniae* (Martinez-Martinez et al., 1998), has 81 Chi sites in its 186 kbp of DNA, or 436/Mbp, twice the density of Chi in *E. coli*’s chromosome. Plasmid pJP4, isolated from *Alcaligenes eutrophus,* even more distantly related to *E. coli* and conferring herbicide-resistance (Don and Pemberton, 1981), has 410 Chi sites/Mbp. Both plasmids move between a wide range of bacteria and belong to different incompatibility groups. So, *E. coli* does not treat plasmids as foreign or not depending upon their Chi content.

Parallel to the wide distribution of Chi, RecBCD or its analog AddAB is encoded by >90% of bacterial species with sequenced genomes (Cromie, 2009). In most of the cases tested, these enzymes are important for DNA repair or recombination or both. Below, we discuss the dogma that Chi or other short sequences prompt these enzymes to promote recombination or enable them to distinguish self *vs.* non-self DNA.

### Chi is active in “foreign” species

If Chi allows *E. coli* to tell self from non-self DNA, one would expect Chi to be active only in *E. coli*. This is not the case. Extracts of other enteric bacteria, including *K. pneumoniae,* and even more distantly related *Proteus mirabilis* and *Vibrio harveyi* nick DNA in a Chi-specific manner, as do *E. coli* extracts or purified RecBCD (Table 2) (Schultz and Smith, 1986) (see Figure 1 and section below on a myth about phage destruction for discussion of RecBCD’s action on DNA with Chi). When the *recBCD* genes of these enteric bacteria are expressed in *E. coli*, they confer Chi hotspot activity in genetic recombination assays and Chi-dependent nicking activity in enzymatic assays (McKittrick and Smith, 1989). Chi is also active in lambda crosses conducted in *Salmonella enterica* cells (Smith et al., 1986). Thus, species other than *E. coli* have high density Chi sites, and they activate them as recombination hotspots. Clearly, Chi is not restricted to *E. coli*.

**Table 2.** Chi is active beyond E. coli, in all tested enteric bacteria.

| Species | Chi nicking* | Chi hotspot activity** |
| --- | --- | --- |
| *Escherichia coli* | +++ | 7.3 |
| *Salmonella enterica* | +++ | 3.7*** |
| *Klebsiella pneumoniae* | + | 4.1 |
| *Serratia marcescens* | + | 4.1 |
| *Proteus mirabilis* | ++ | 2.7 |
| *Vibrio harveyi* | +++ | ND |
| *Pseudomonas aeruginosa* | - | 1.0 |
| *Pseudomonas putida* | - | 1.0 |

\* Production of ssDNA cut at Chi during unwinding by RecBCD in extracts of native bacteria or of *E. coli* with cloned *recBCD* from other species (Schultz and Smith, 1986; McKittrick and Smith, 1989).
\*\* Enhancement of recombination in an interval with Chi in lambda crosses in *E. coli* cells with cloned *recBCD* from the indicated species (McKittrick and Smith, 1989). A value of 1 indicates no enhancement (no Chi activity).
\*\*\* 3.2 in *S. enterica* cells (Smith et al., 1986).

By contrast, strong evidence indicates that Chi is not active in *Pseudomonas* spp. Extracts of *P. aeruginosa* and *P. putida* do not nick at Chi, unlike extracts of all enteric bacteria tested (Schultz and Smith, 1986; McKittrick and Smith, 1989). Chi is not active as a recombination hotspot in *E. coli* cells expressing *recBCD* genes from either *Pseudomonas* species, but it is active in cells expressing each of the other enteric RecBCD enzymes tested (McKittrick and Smith, 1989). RecBCD enzyme purified from *P. syringae* cuts DNA at a different, but closely related, sequence 5’ GCTGGCGC 3’ (Pavankumar et al., 2018). This sequence may be a recombination hotspot in *P. syringae*, but to our knowledge this has not been directly demonstrated. If it indeed is a recombination hotspot, its sequence similarity to that of *E. coli* Chi provides excellent opportunity for exploring the evolution of hotspots.

### Is Chi “over-represented” in *E. coli*’s genome?

It is often stated, as part of the dogma discussed above, that Chi occurs in the *E. coli* genome more frequently than expected. For example, “Since Chi sites occur every ∼5 kb in the *E. coli* genome, which is about 14 times more frequent than expected by chance, these sites were suggested as markers of bacterial self, … [and are] responsible for some of the self/non-self discrimination properties of the CRISPR adaptation process” (Levy et al., 2015). The basis of this expectation is not stated, but presumably it assumes random association of nucleotides in the *E. coli* genome. Of course, the nucleotide composition of compact bacterial genomes is rarely random.

We therefore analyzed the *E. coli* genome for features that might account for Chi’s density in *E. coli*. For the first analyses, we used only the “top” strand, that with 5’ AGCTTTTCATTCTG 3’ at nucleotides 1 – 14 in NC_000913.3, the sequence of *E. coli* strain K12, substrain MG1655. In this way, we might see asymmetries or other relations that would be missed using both strands. (Analysis of the bottom strand or both strands together gives similar results.) Table S2 gives for each nucleotide its count and frequency, which is close to, but not equal to, 0.25 for each nucleotide. Taking the observed frequencies and assuming random association, we calculate 149.9 Chi sites/9.28 million nucleotides (counting both strands, or 149.9/4.64 Mbp, in the *E. coli* genome). This is 6.72 times less than observed (1008/4.64 Mbp), as we noted previously (Colbert et al., 1998). (The value of 14 times quoted above may not have counted the nucleotides of both strands.) Thus, Chi is present more frequently than random nucleotide association predicts, leaving open the question of what, if anything, accounts for Chi’s high frequency in *E. coli* DNA.

### Nucleotides are not randomly arranged in the *E. coli* genome

We tested the randomness of nucleotides in the *E. coli* genome in various ways and observed high non-randomness. If nucleotides were randomly associated, the frequency of each dinucleotide (TT, TA, AT, *etc.*) should be the product of the individual nucleotide frequencies (Table S2). These calculated values are far from the observed values (Figure 2A); counting dinucleotides as non-overlapping (1 2, 3 4, 5 6, *etc.*) or as overlapping (1 2, 2 3, 3 4, *etc.*) gave indistinguishable results (Figure S1). Thus, neighboring nucleotides are highly correlated. But there is no significant correlation of nucleotides 10000 apart, for example (Figure 2B). We expected this result from consideration of neighboring nucleotides being correlated due to non-randomness of codon and amino acid frequencies, protein-coding sequences (open reading frames, or ORFs) being less than 10000 nucleotides long (Figure 3), and ORFs occupying 87.5% of *E. coli*’s genome. Beyond ORF lengths, we expected no nucleotide correlation, as observed. We then explored ORF features that might account for the observed frequency of Chi in the *E. coli* genome.

**Figure 2.**
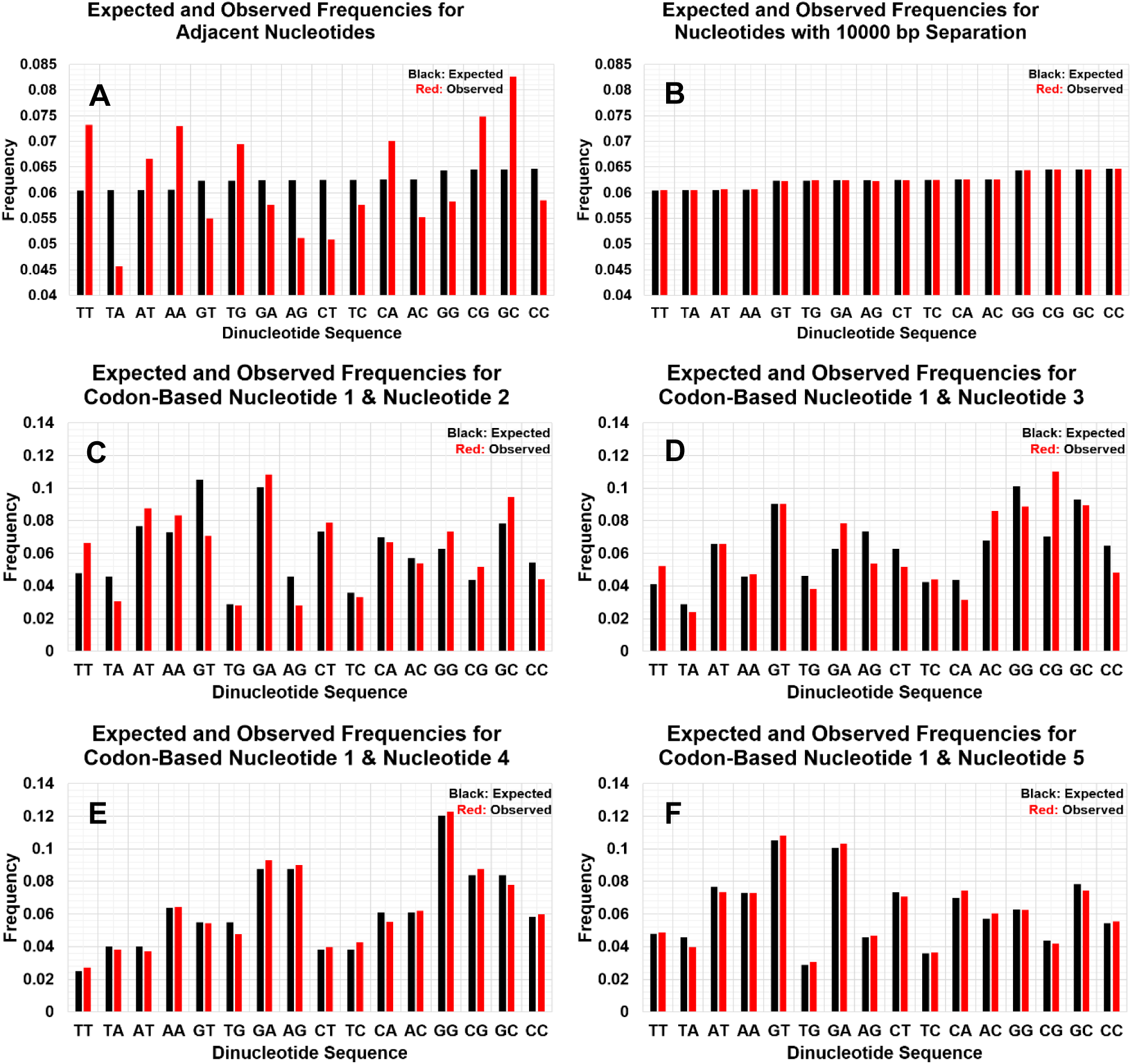
Frequencies of close nucleotide pairs are not random, depend on the nucleotide positions within codons, and display a repeating pattern. Note differences in vertical scales. (**A** and **B**) The expected frequencies of each nucleotide pair, the product of the individual nucleotide frequencies in the *entire top strand* of *E. coli* (Table S2), are in black; in red are the observed frequencies for adjacent nucleotides (0 separation) (**A**) and for nucleotides separated by 10000 nucleotides (**B**). (**C – F**) The expected frequencies of each nucleotide pair, the product of the individual nucleotide frequencies at *each codon position* in all ORFs (Table 3), are shown for adjacent nucleotides (0 separation) (**C**) and for nucleotides separated by 1 (**D**), 2 (**E**), or 3 (**F**) nucleotides in black. Corresponding observed nucleotide pair frequencies are in red. Note that separation by 0, 1, or 2 nucleotides have different patterns, both expected and observed, but separation by 3 nucleotides (nucleotides in positions 1 and 5; **F**) resembles that of 0 nucleotide separation (nucleotides in positions 1 and 2; **C**).

**Figure 3.**
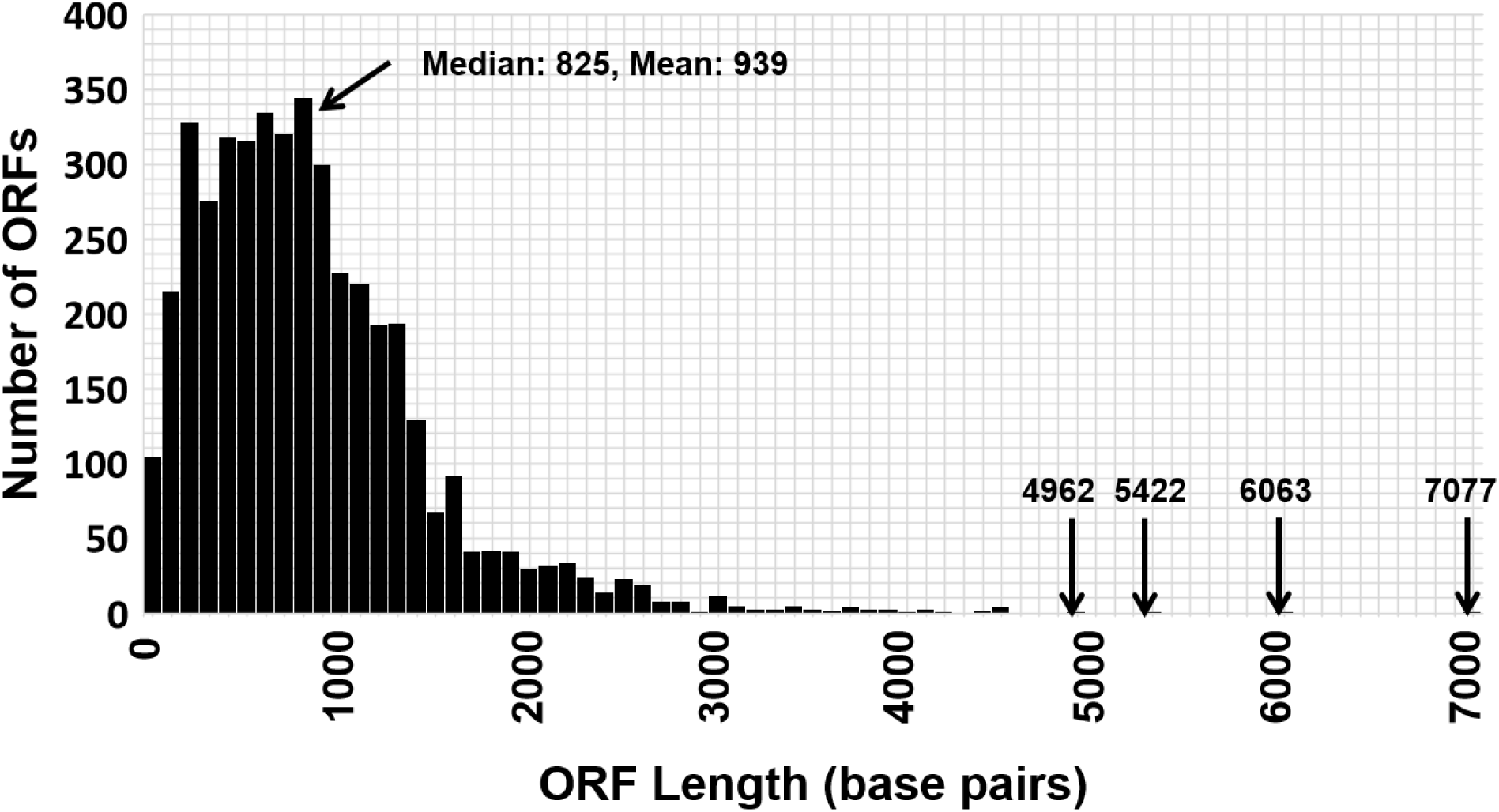
The distribution of *E. coli* ORF lengths accounts for nucleotide associations becoming nearly random between about 1500 and 3500 bp. Shown are the number of ORFs (vertical axis) in each size range (bins of 100 bp; horizontal axis) and the four individual ORFs >4612 bp long. Figures 2 and S3 – S6 show that the non-randomness of adjacent nucleotides is greater than that of nucleotides separated by many nucleotides; the non-randomness becomes progressively less at 751, 1501, 3502, and 10000 nucleotides of separation. This distance-dependence is accounted for by most ORFs being less than 1500 nucleotides long.

**Table 3.** Chi contains *E. coli*’s most frequent codon 5’ CTG 3’.

| Codon | Number | Codon | Number | Codon | Number | Codon | Number |
| --- | --- | --- | --- | --- | --- | --- | --- |
| †(F)TTT | 30335 | (S) TCT | 11527 | (Y) TAT | 22013 | (C) TGT | 7064 |
| (F) TTC | 22534 | (S) TCC | 11755 | (Y) TAC | 16670 | (C) TGC | 8871 |
| (L) TTA | 18957 | (S) TCA | 9780 | (*) TAA | 2875 | (*) TGA | 1416 |
| (L) TTG | 18659 | (S) TCG | 12209 | (*) TAG | 359 | (W) **TGG** | 20896 |
| (L) CTT | 15020 | (P) CCT | 9544 | (H) CAT | 17610 | (R) CGT | 28514 |
| (L) CTC | 15119 | (P) CCC | 7468 | (H) CAC | 13232 | (R) CGC | 30013 |
| (L) CTA | 5325 | (P) CCA | 11545 | (Q) CAA | 20957 | (R) CGA | 4885 |
| (L) **CTG** | 71938 | (P) CCG | 31705 | (Q) CAG | 39361 | (R) CGG | 7408 |
| (I) ATT | 41473 | (T) ACT | 12179 | (N) AAT | 24141 | (S) AGT | 11958 |
| (I) ATC | 34324 | (T) ACC | 31879 | (N) AAC | 29466 | (S) AGC | 21896 |
| (I) ATA | 5952 | (T) ACA | 9641 | (K) AAA | 45902 | (R) AGA | 2818 |
| (M) ATG | 37931 | (T) ACG | 19701 | (K) AAG | 14037 | (R) AGG | 1612 |
| (V) GTT | 24933 | (A) **GCT** | 20794 | (D) GAT | 43790 | (G) **GGT** | 33733 |
| (V) GTC | 20833 | (A) GCC | 34784 | (D) GAC | 26037 | (G) GGC | 40443 |
| (V) GTA | 14843 | (A) GCA | 27535 | (E) GAA | 53882 | (G) GGA | 10800 |
| (V) **GTG** | 35736 | (A) GCG | 45941 | (E) GAG | 24261 | (G) GGG | 15091 |
| Total Codons: 1363910 [4650 ORFs] |  |  |  |  |  |  |  |

† Letters in parentheses indicate the encoded amino acid, using the one-letter abbreviations in Table S3.
Data are for all ORFs ≥14 codons long as annotated in *E. coli* strain K-12 substrain MG1655 (GenBank accession number NC\_000913.3). Bold-face codons are in Chi (5'GCTGGTGG 3').

### Amino acids are not quite randomly associated in *E. coli* proteins

The frequency of amino acids encoded by *E. coli* ORFs ranges from 10.7% for Leu to 1.2% for Cys (Table S3). The observed frequency of each dipeptide (LeuAla, ArgTrp, *etc.*) is similar to, but not exactly, that expected from random association (Figure 4). Pearson’s correlation coefficient is 0.98, but the slope of the linear regression line (0.94 ± 0.01) is highly significantly different from 1, the value that indicates random association (p < 0.001). Certain amino acid pairs occur noticeably more, and some less, frequently than expected from random association. For example, the observed and expected frequencies for ThrLeu are 0.723% and 0.572%; those for ValTyr are 0.200% and 0.324% (p < 0.001 by two-tailed chi-square test). These results led us to examine codons for non-randomness.

**Figure 4.**
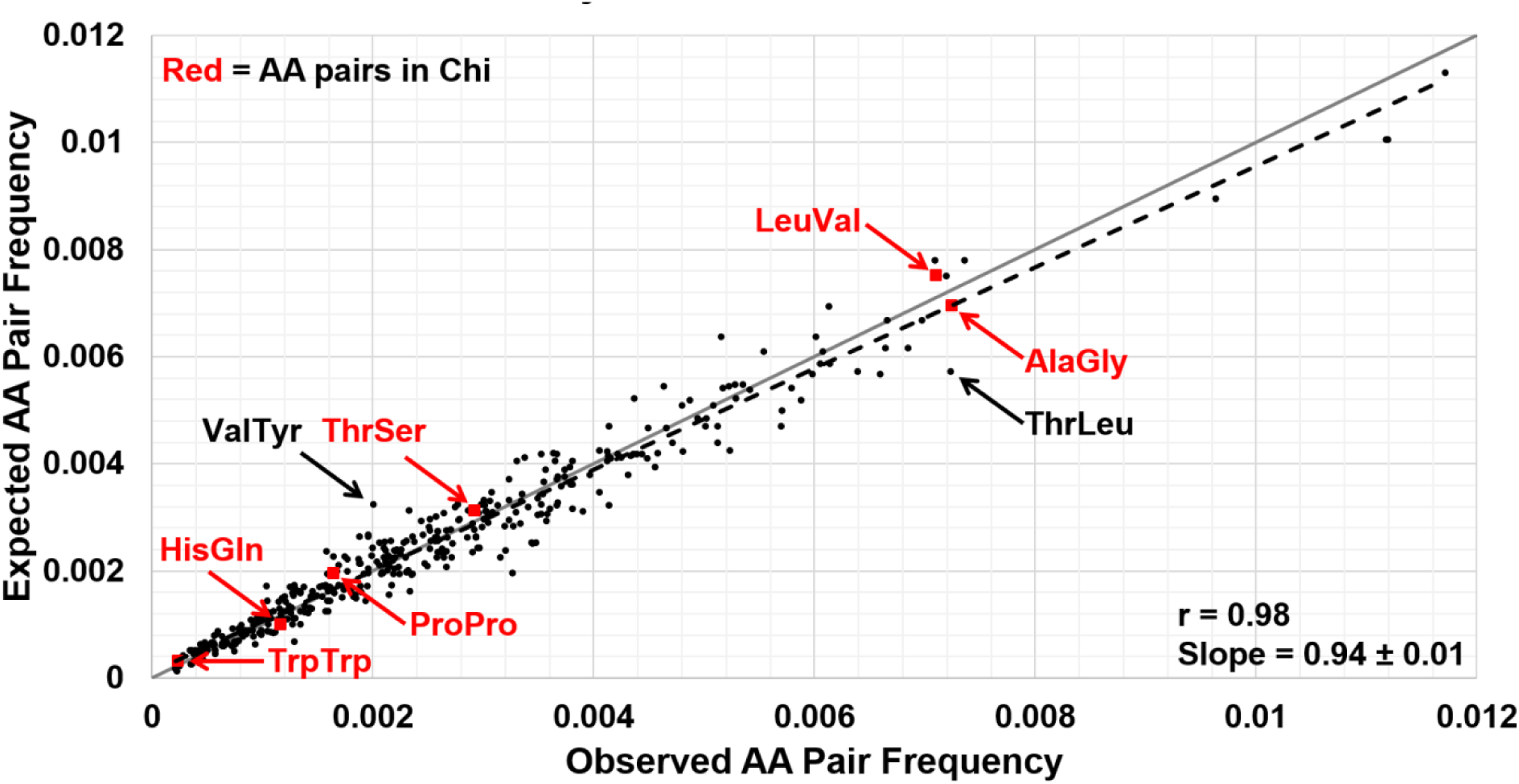
*E. coli* amino acids are not quite randomly associated. The frequency of each pair of adjacent amino acids (AA) in all ORFs is plotted against its randomly expected frequency (product of individual amino acid frequencies in Table S3). Note that the slope of the linear regression line (dashed black) is significantly different from 1 (solid grey line, p < 0.0001 by two-sided unpaired t test), which would indicate randomness; Pearson correlation coefficient **r** is near unity, indicating high correlation. In red are dipeptides encoded by Chi and its complement. In black are two dipeptides whose observed frequencies are especially different from their expected frequencies (p < 0.0001 by two-tailed chi-square test). See also Figure S2.

### Codon usage is even more non-random in *E. coli*

Codon usage, like amino acid usage, varies highly. The most frequent codon, CTG for Leu, is used 45 times more frequently than the least frequent codon, AGG for Arg, even though both are abundant amino acids (Tables 3 and S3). Even among codons for Leu, CTG is 14 times more frequent that CTA. (Note that CTG is part of Chi.) Codons are also non-randomly associated: the observed frequency of each codon pair is markedly different from their random association (Figure 5), as noted by others (Gutman and Hatfield, 1989; Boycheva et al., 2003) . (For calculations involving codons, we used all ORFs with ≥14 codons on both strands. ORFs on opposite strands only rarely overlap, providing non-redundant data.) Pearson’s correlation coefficient is 0.71, and the slope of the linear regression line (0.77 ± 0.01) indicates high non-randomness. These not-surprising results led us to seek some feature of ORFs and codon usage that might account for the observed frequency of Chi.

**Figure 5.**
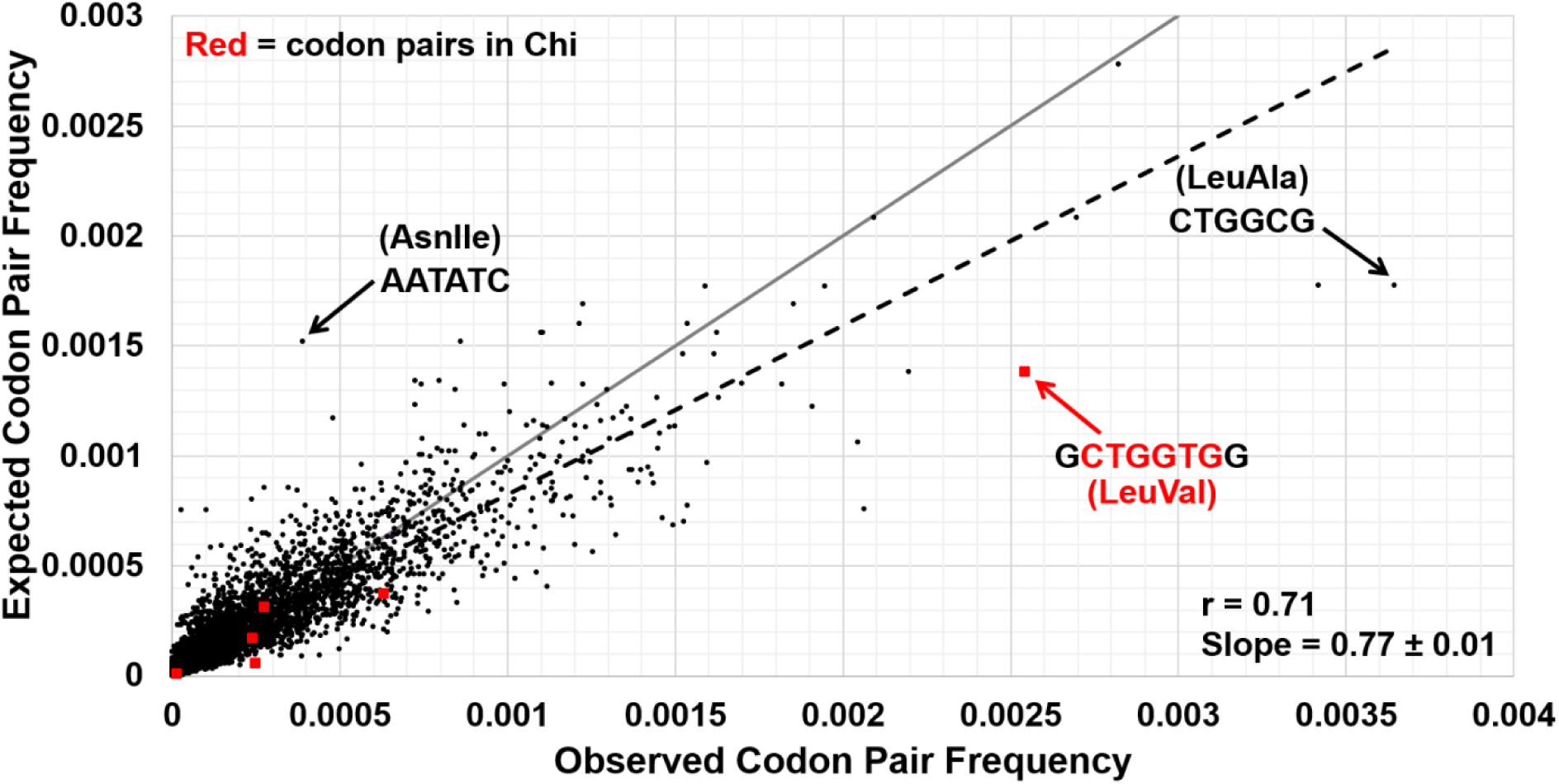
*E. coli* codons are not randomly associated. The frequency of each pair of adjacent codons in all ORFs is plotted against its expected frequency (product of individual codon frequencies in Table 3). Note that the slope of the linear regression line (dashed black) is significantly different from 1 (solid grey line, p < 0.0001 by two-sided unpaired t test), which would indicate randomness; Pearson correlation coefficient **r** (0.71) also indicates significant but incomplete correlation. In red are dicodons contained in Chi and its complement. In black are two dicodons whose observed frequencies are especially different from their expected frequencies (p < 0.0001 by unpaired two-tailed chi-square test). See also Figure S2.

We analyzed amino acids separated by one amino acid and found that these pairs are more nearly randomly associated than adjacent amino acids (0 separation) (Figures S2A and S2C); pairs separated by two amino acids are even more nearly random. Similar relations occur for adjacent codons and those separated by one or two codons (Figure S2B and S2D). These results show, as expected, progressively less correlation as amino acids or codons are farther apart and reinforce the non-randomness of the *E. coli* genome.

### Preferential codon usage can account for Chi’s high frequency in *E. coli*

We next asked whether the frequency of codon usage could account for the non-randomness of dinucleotides noted above. Although adjacent codons are not randomly associated (Figure 5), we assumed randomness as a means to obtain “expected” values for comparison with the observed values in the following considerations. [ORFs occupy 42.5% of the top strand and 45.0% of the bottom strand; 99% (997/1008) of Chi sites are in ORFs. Thus, these calculations consider the genome portion containing nearly all Chi sites.] First, we calculated the frequency of dinucleotides expected from random association of codons. For this, we determined the frequency of each nucleotide at each position in codons (Table 4). We then calculated the frequency of each dinucleotide (TT, TA, AT *etc.*) at positions 1&2, 1&3, 1&4, …, 2&3, 2&4, 2&5, …, and 3&4, 3&5, 3&6, *etc.* expected from random nucleotide association and compared these with the observed frequencies (Figures 2C-F). (We use “&” between nucleotides, *e.g*. 1&2 in Figures 2C-F, when their positions in codons are considered and a space, *e.g*. 1 2 in Figures 2A-B, when codon position is not considered.)

**Table 4.**
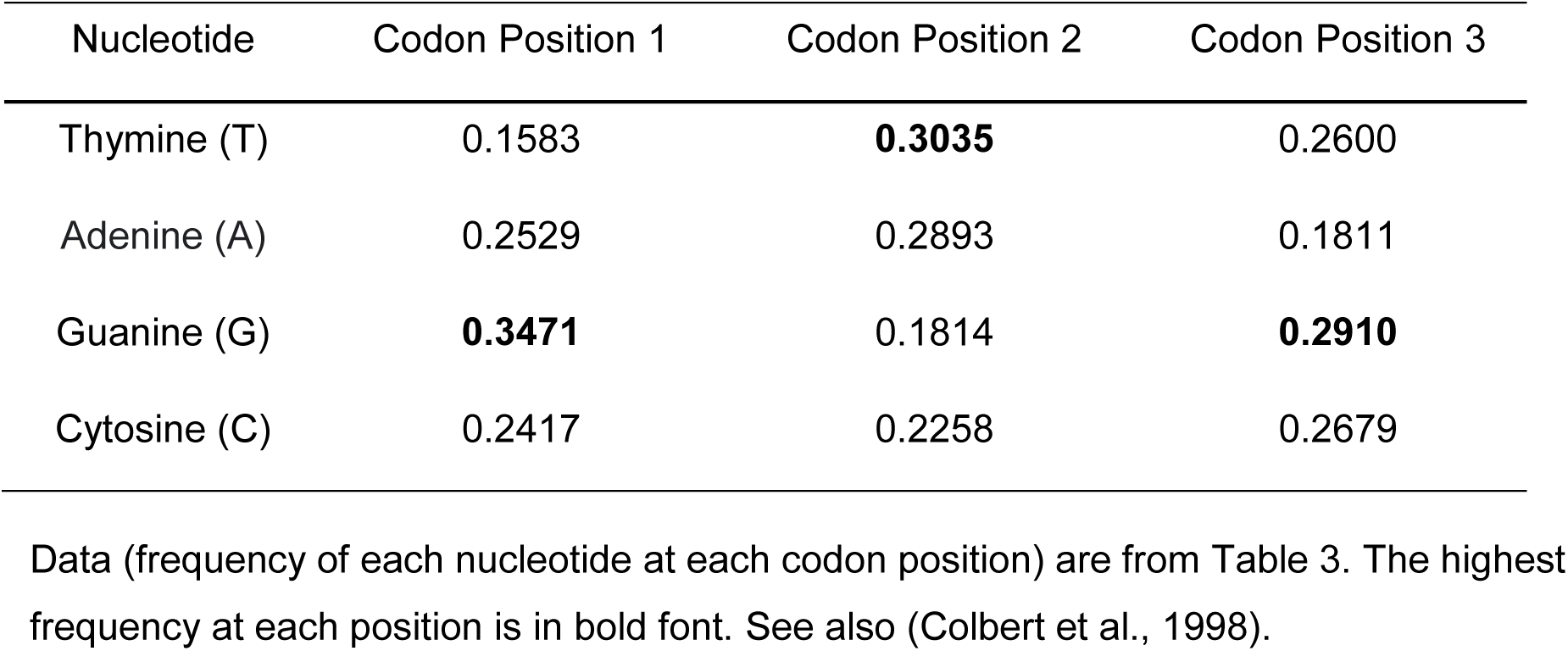
Chi contains E. coli’s “consensus” codon 5’ GTG 3’.

| Nucleotide | Codon Position 1 | Codon Position 2 | Codon Position 3 |
| --- | --- | --- | --- |
| Thymine (T) | 0.1583 | **0.3035** | 0.2600 |
| Adenine (A) | 0.2529 | 0.2893 | 0.1811 |
| Guanine (G) | **0.3471** | 0.1814 | **0.2910** |
| Cytosine (C) | 0.2417 | 0.2258 | 0.2679 |

Consider first dinucleotides composed of nucleotides at the first and second positions (1&2) in each codon. The observed frequencies of TT, TA, AT *etc.* are similar, but not identical, to the expected frequencies (Figure 2C). Thus, codon usage accounts for much of the non-randomness of nucleotides in the *E. coli* genome. Note that the range of observed frequencies considering ORFs and codon usage (from 0.0280 for TG to 0.108 for GA; Figure 2C) is greater than the range observed (from 0.0457 for TA to 0.0828 for GC) for all dinucleotides without ORF or codon consideration; Figure 2A). This codon-based analysis reveals an even stronger non-random feature of *E. coli*’s genome than the simple nucleotide-based analysis.

We similarly considered dinucleotides composed of the first and third positions (1&3) in each codon. Again, the observed frequencies in most cases are similar to the expected (Figure 2D), but the pattern differs from that of positions 1&2 (Figure 2C). Dinucleotides at positions 1&4 (*i.e.*, the first position of adjacent codons; Figure 2E) have a frequency distribution that differs even more than those of 1&2 or 1&3. The frequency distribution of 1&5 (Figure 2F), however, is similar to that of 1&2, as expected if codon, and thus nucleotide, association becomes more nearly random with increasing distance. We quantified these comparisons (Figure S3) and found, as expected, that the correlations between observed and expected frequencies are higher for identities (*e.g.,* observed 1&2 compared with expected 1&2) (Figure S3A) than for non-identities (*e.g.*, 1&2 compared with 1&3) (Figure S3B) except for 1&2 compared with 1&5 (Figure S3C). These results confirm that codon usage accounts for much of *E. coli*’s non-random genome structure.

Going back to examination of nucleotide pairs on the top strand without consideration of ORFs or codons (Figures 2A and B), we noted that the non-random pattern noted above persisted for quite some distance between nucleotides in the pair but became less noticeable in the range of several hundred bp (Figures S4 and S5). There was, however, a 3 bp periodicity in the observed patterns, as expected from a major role for codon usage in determining the non-randomness discussed here. First, consider pairs of nucleotides separated by 2 or 5 nucleotides (*e.g.*, nucleotides 1 4 or 1 7), which would be in the same codon positions (*i.e.*, first, second or third position) if they are in the same ORF (there are no known introns in *E. coli* genes). The most frequent dinucleotides separated by 2 nucleotides are also the most frequent dinucleotides separated by 5 nucleotides; similarly for the least frequent dinucleotides, although in each comparison the magnitudes differ (Figure S5A and D). Thus, the observed patterns of separation by 2 or 5 nucleotides are related. Furthermore, separations by 3 or 6 nucleotides are similar (Figure S5B and E), as are separations by 4 or 7 nucleotides (Figure S5C and F). At 749 – 754 nucleotide separations, the dinucleotide frequency distributions are still non-random but not as highly non-random as with less separation (*e.g.*, 0, 2, 3, or 4 nucleotides; Figures 2A and S5). The pattern with separation of 749 nucleotides (*e.g.,* 1 751) is similar to that with separation of 752 nucleotides (*e.g*., 1 754), just as 1 4 is similar to 1 7, *etc.* (Figures S5A and D). These 3-nucleotide periodicities are expected from the influence of codon usage. The pattern becomes more nearly like that of random nucleotide association (Figure 2A) as the separation increases to 1501 and 3502 nucleotides (Figures S4 and S6). With 10000 nucleotides of separation (Figure 2B), the pattern is indistinguishable from that of random association, as noted before. These outcomes are expected from the distribution of *E. coli* ORF lengths (Figure 3).

These analyses show that the non-randomness of *E. coli*’s genome stems from codon usage. They support the view that the often-stated “unexpectedly” or “disproportionately” high frequency of Chi is due to its codon usage, as noted by others and one of us (Biaudet et al., 1998; Colbert et al., 1998; Uno et al., 2006), rather than recombination-based selection enriching this octamer. We redid the calculations by Colbert et al. (Colbert et al., 1998) for the expected and observed frequency of Chi in each reading frame (Table 5) using a slightly different genome sequence and set of ORFs for codon usage. The results confirm that Chi occurs most often with CTG in the reading frame, as expected from CTG being the most frequent codon for the most frequent amino acid (Leu) (Tables 3 and S3). The expected frequencies of Chi or its complement (5’ CCACCAGC 3’) in each reading frame are similar to the observed frequencies. Thus, the observed frequency of Chi can be nearly fully accounted for by its sequence, *E. coli*’s codon usage, and ORFs occupying most of the genome.

**Table 5.**
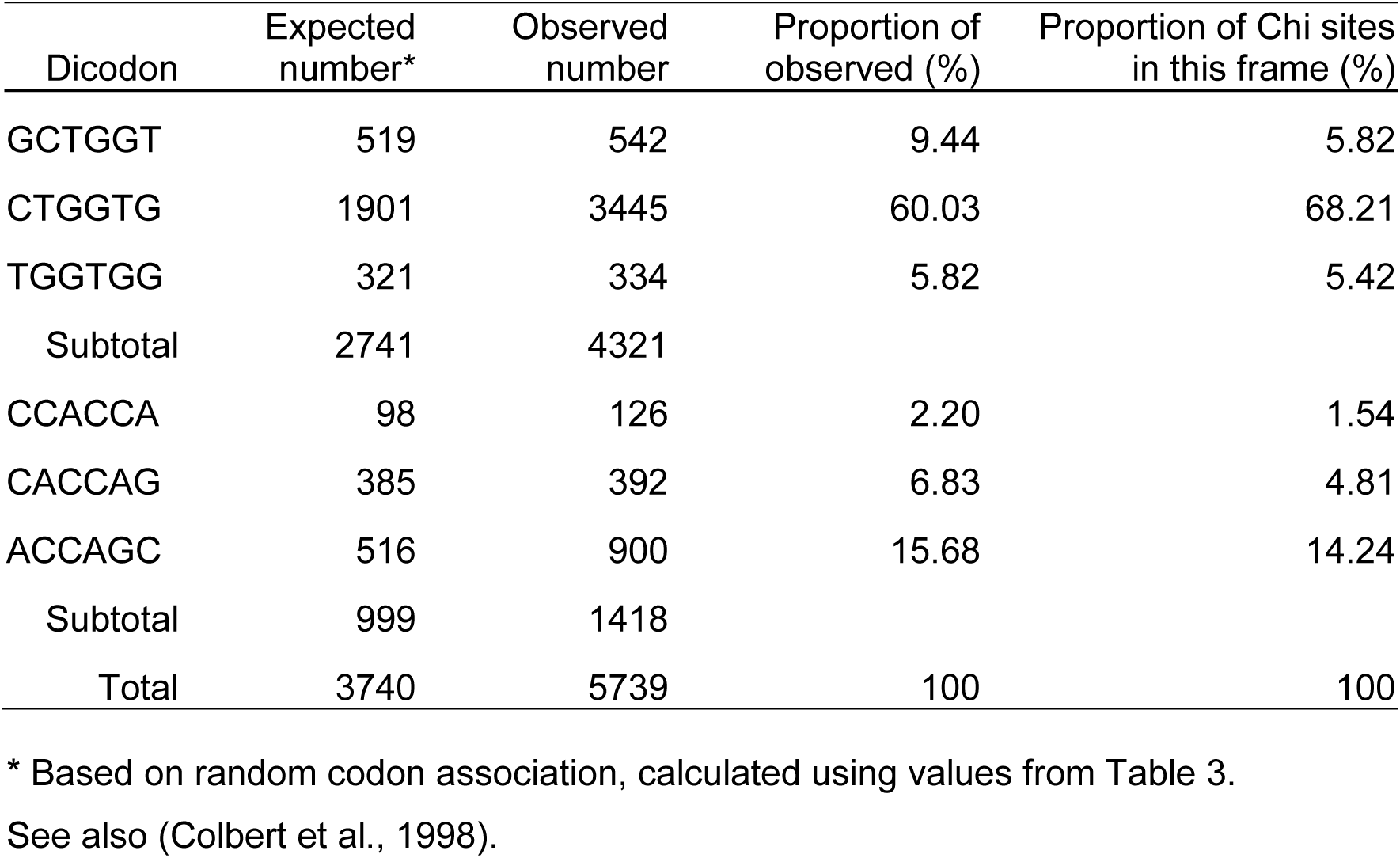
Chi’s frequency is correlated with its in-frame dicodon frequencies.

| Dicodon | Expected number* | Observed number | Proportion of observed (%) | Proportion of Chi sites in this frame (%) |
| --- | --- | --- | --- | --- |
| GCTGGT | 519 | 542 | 9.44 | 5.82 |
| CTGGTG | 1901 | 3445 | 60.03 | 68.21 |
| TGGTGG | 321 | 334 | 5.82 | 5.42 |
| Subtotal | 2741 | 4321 |  |  |
| CCACCA | 98 | 126 | 2.20 | 1.54 |
| CACCAG | 385 | 392 | 6.83 | 4.81 |
| ACCAGC | 516 | 900 | 15.68 | 14.24 |
| Subtotal | 999 | 1418 |  |  |
| Total | 3740 | 5739 | 100 | 100 |

\* Based on random codon association, calculated using values from Table 3.
See also (Colbert et al., 1998).

Chi’s non-random orientation on the chromosome can also be accounted for by *E. coli*’s non-random orientation of ORFs (Bell et al., 1998; Colbert et al., 1998). Chi is oriented on the *E. coli* genome in a manner consistent with Chi being active in repair of DNA broken during replication. When a replication fork is broken on either the leading or lagging strand, RecBCD can enter the two ds DNA ends and promote restoration of the fork (Kuzminov et al., 1994). Entry onto the end heading RecBCD toward the origin of replication, but not onto the other end, would lead to the entered end being rejoined to the intact DNA homologous to it and allow replication to restart at the new fork. Chi oriented with the entry end to the right of 5’ GCTGGTGG 3’ would activate RecBCD when it encounters Chi (Taylor et al., 1985). 75.5% of Chi sites are oriented in this way (replication proceeds in both directions from the unique origin). Genes, especially highly expressed, essential ones, are preferentially oriented such that they are transcribed, and accordingly translated, in the same direction as replication, presumably to avoid head-on collisions of transcription and replication (Brewer, 1988; Rocha, 2004; Schroeder et al., 2020). These biases thus orient Chi in the direction favoring repair of broken replication forks. It follows that the biased orientation of Chi is no mystery – it stems from transcription, and consequently translation, being biased in the direction of replication plus non-random codon usage.

Buton and Bobay (2021) attempted to predict the Chi-equivalent sequence in other Proteobacteria, based on assumptions of similarity to Chi’s sequence, high frequency, and chromosomal orientation-dependence. They failed to detect the “Chi” sequence of *H. influenzae* noted below, suggesting that these computational methods are inadequate. We suspect only experimental searches will successfully identify recombination hotspots.

What led to Chi being 5’ GCTGGTGG 3’? It is clearly highly represented in the genome – Chi and its complement are the 21^st^ and 22^nd^ most common octamers of 65,484 present on the two strands of *E. coli* DNA (Figure 6). The “consensus codon” GTG, that with the most frequent nucleotide at each position (Table 4), and CTG, the most frequent codon (Table 3), are both in Chi. 5’ CTG|GTG 3’, the central part of Chi, contains the most frequent codons for Leu and Val, two of the four most-frequent amino acids encoded by *E. coli* (Tables 3 and S3) and is the fifth most abundant dicodon (Figure 5). These features likely have something to do with 5’ GCTGGTGG 3’ being “chosen” as Chi. But these are features of translation, not of recombination. We suppose that an ancient ancestor of *E. coli* had a RecBCD-like DNA repair enzyme with no sequence preference. A *recBCD* mutant with weak preference for some DNA sequence arose and had a higher efficiency of repair. Additional mutations with increasing efficiency gained abundance in the population. One that used a highly frequent sequence, such as 5’ GCTGGTGG 3’, would have been the most efficient and abundant. Perhaps *E. coli* is still struggling to recognize the most frequent octamer 5’ C<u>GCTGG</u>C<u>G</u> 3’ in its genome.

**Figure 6.**
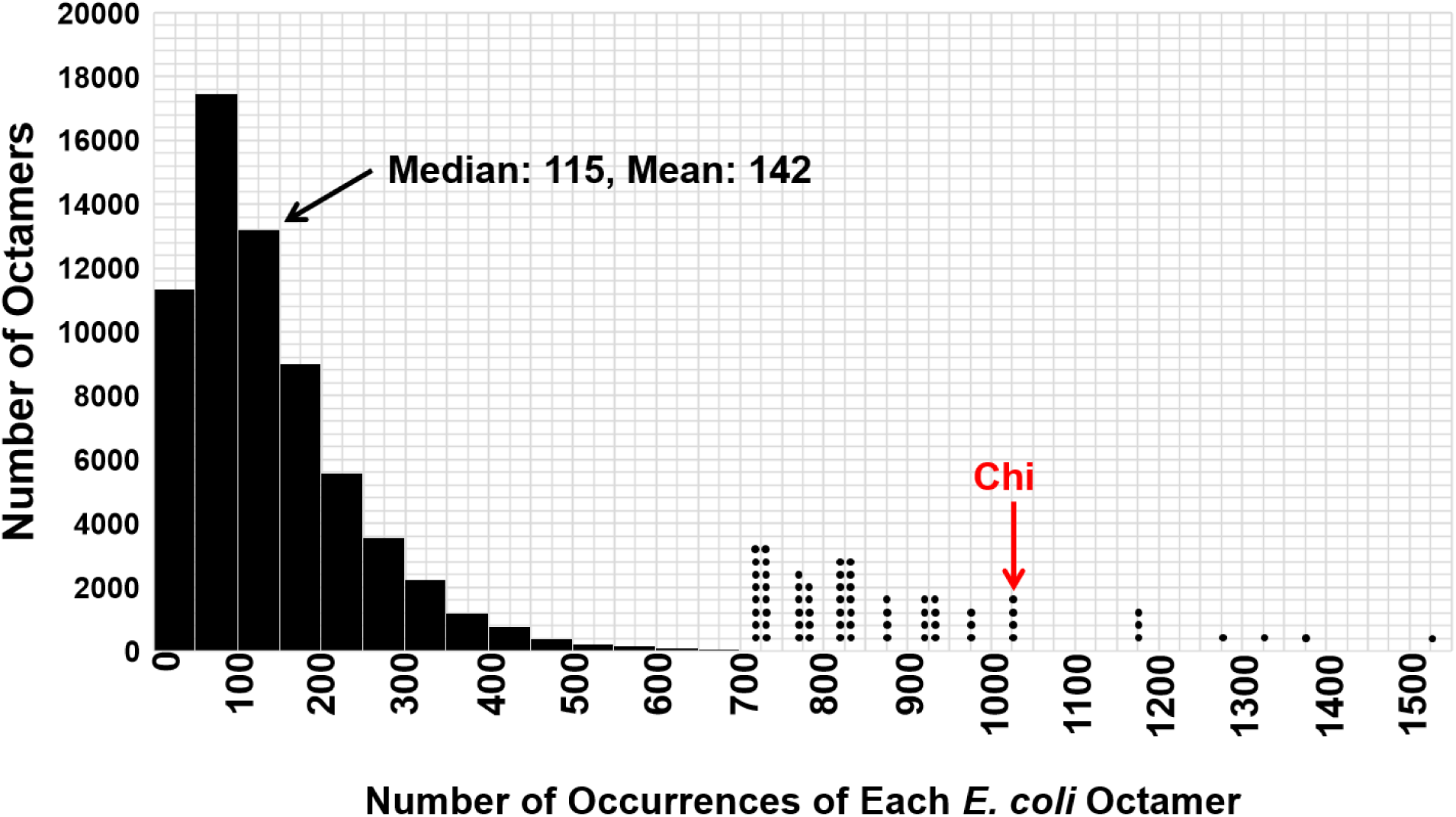
Chi is one of the most frequent *E. coli* octamers. The number of occurrences in the *E. coli* genome (both strands) of each octamer-complement pair, in bins of 50 (horizontal axis), is plotted against the number of octamer-complement pairs in that frequency range (vertical axis). There are 65484 octamers in the genome. Those occurring >700 times each are shown by individual bullets (●), one for each octamer-complement pair. Chi and its complement (red arrow) occur 1008 times.

### “Chi” sequence of *S. aureus* also comports with its codon usage

The view developed here can also account for the sequence 5’ GAAGCGG 3’ presumably affecting the activity of the *Staphylococcus aureus* AddAB enzyme, its RecBCD analog (Halpern et al., 2007). This sequence specificity was identified from studies of rolling circle replication. Some phages, including lambda, and some plasmids change from circular (theta, θ) replication to rolling circle (sigma, σ) replication during their infection cycles. The end of the rolling circle can be degraded by RecBCD or AddAB nuclease activity. For example, *S. aureus* plasmid pC194 replicating as a rolling circle in *E. coli* cells forms high molecular weight (HMW) DNA only if the plasmid contains Chi, providing a convenient screen for DNA containing a special sequence (Dabert et al., 1992). In *S. aureus*, this plasmid makes HMW DNA if it contains 5’ GAAGCGG 3’ (Halpern et al., 2007). Thus, this sequence is often considered to be *S. aureus*’s “Chi” (but see next paragraph). This heptamer and its complement occur 335 times in the *S. aureus* 2.82 Mbp genome and are the 33^rd^ and 34^th^ most frequent heptamers. (By contrast, Chi 5’ GCTGGTGG 3’ ranks 21,806 among *S. aureus*’s 65,327 octamers.) 89.7% of these “Chi” sequences are co-oriented with replication and 92.8% occur in ORFs. *S. aureus*’s codon usage can account for the frequency of “Chi,” just as for *E. coli*’s Chi (Tables S4 and S5). Thus, this sequence shares many properties with Chi and may be a recombination hotspot, but to our knowledge that feature and direct interaction with AddAB have not been demonstrated.

Other sequences that stimulate HMW DNA formation of pC194 have been found for other species and have been called “Chi.” These species and their “Chi” sequences include *B. subtilis* (5’ AGCGG 3’, the 3’ end of the *S. aureus* “Chi” sequence), *Lactococcus lactis* (5’ GCGCGTG 3’), and *Haemophilus influenzae* [5 ’G(G/C)TGGAGG 3’ and 5’ GNTGGTGG 3’, both similar to *E. coli* Chi]. Each of these has been called a hotspot of recombination (Biswas et al., 1995; Chedin et al., 1998; El Karoui et al., 1998; Sourice et al., 1998), but we are unaware of any recombination assays to demonstrate this property. Purified *B. subtilis* AddAB does cut DNA at its “Chi” site, showing, as expected, a direct interaction with this recombination-promoting enzyme (Chedin et al., 2000). To our knowledge, a direct interaction between these “Chi” sequences and *L. lactis* AddAB or *H. influenzae* RecBCD has not been reported. Furthermore, *H. influenzae* “Chi” activity in HMW formation is not orientation-dependent, unlike the other Chi and “Chi” sequences. The formation of HMW DNA is often interpreted as “Chi” blocking the nuclease activity of the AddAB or RecBCD enzyme (Dabert et al., 1992; Chedin et al., 1998; Sourice et al., 1998; Halpern et al., 2007). But this simple view does not explain why formation of HMW DNA requires RecA protein (see section below about RecBCD destroying DNA).

### Temperate phage P1 contains 50 Chi sites, likely uses them to its advantage, and appears to select for Chi as a recombination hotspot

We noted above that phage P1 contains Chi at more than twice the density of *E. coli* (Table 1). P1 has 50 Chi sites across its 94.8 kbp genome (Figure 7). These Chi sites are abundant on both strands (31 are on the “top” strand; Table S6), but their positions are clearly non-random. Chi is much denser toward the “ends” of the genome than in the middle of the genome. We understand this non-random distribution as follows. During its lytic phase, P1 replicates as concatemers about four units long (Sternberg and Hoess, 1983). It initiates packaging by cutting the DNA at its unique *pac* site and proceeding in one direction (to the right as shown in Figure 7). When it reaches the intact *pac* site one genome length away, it proceeds ∼5 – 10 kbp farther and cuts again (Ikeda and Tomizawa, 1968; Yun and Vapnek, 1977; Iida and Arber, 1979). This linear DNA, with terminal repeats, is packaged into the phage particle (virion). The next phage DNA is about the same length as the first and has ∼5 – 10 kbp terminal repeats of the genome sequence adjacent to the first terminal repeats. Packaging proceeds similarly for about four units. Thus, the far end of the fourth packaged DNA is ∼20 – 40 kbp from the initial *pac* site. Most of the phage DNA ends are thus in genomic regions with dense Chi sites (Figure 7).

**Figure 7.**
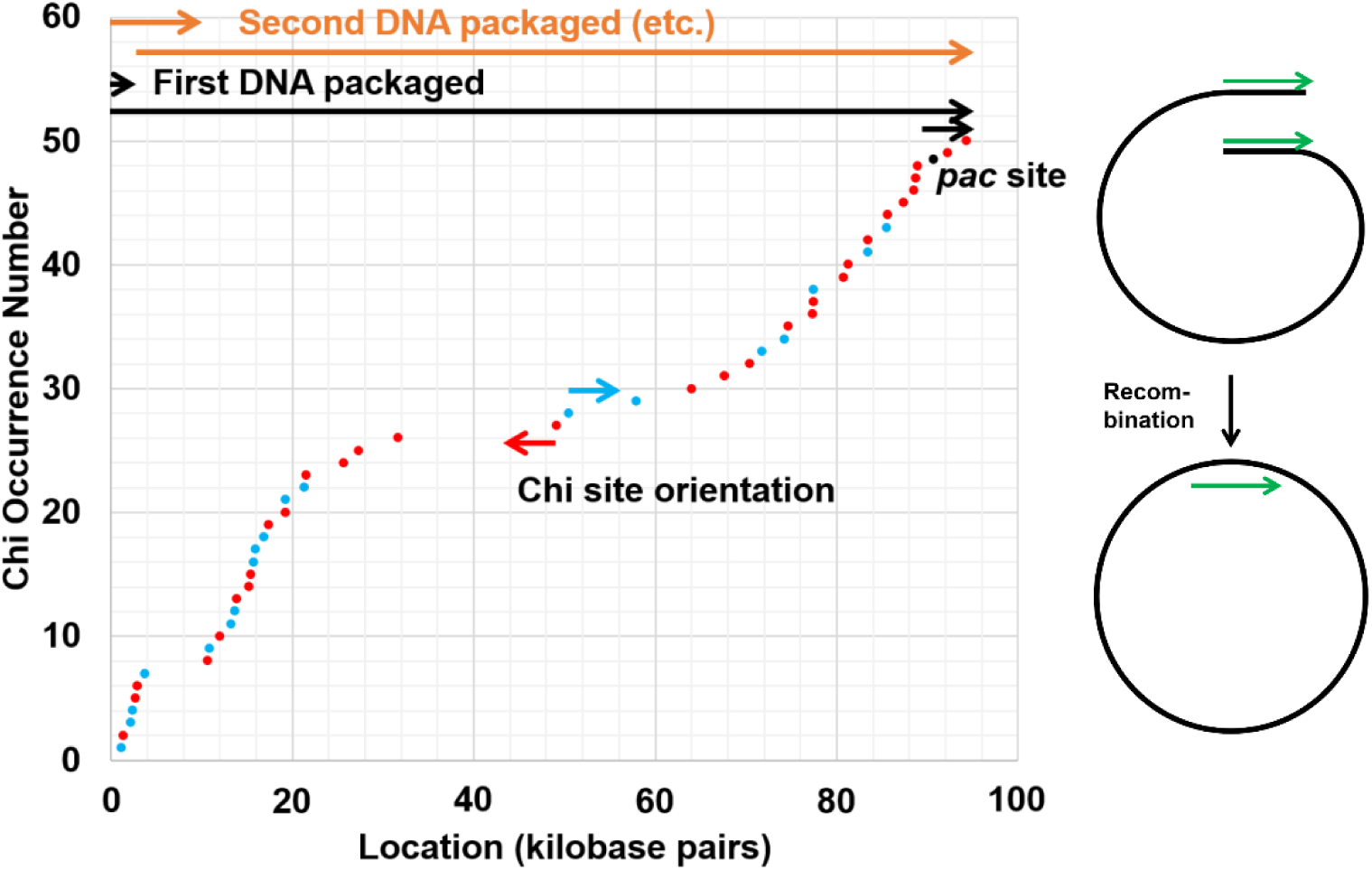
High Chi-density in *E. coli* phage P1 may reflect the need for RecBCD to circularize its injected linear DNA. Chi sites are placed according to their numerical order from the left end on the unique P1 genome sequence. Chi sites at red dots activate RecBCD unwinding DNA from a ds end to its right; those at blue dots activate RecBCD from the left. Note the higher Chi density toward each end. See text for description of sequential DNA packaging beginning at the unique *pac* site (black dot). Diagram on the right illustrates recombination between the terminal direct repeats (∼5 – 10 kb; green arrows) to form circular DNA essential both for lysogenic propagation as a plasmid and for lytic growth to produce concatemers for packaging into virions. We propose that P1 and P7 use Chi sites to the right of the *pac* site, as shown here, and D6 and NGUA22, with inverted *pac* sites, use Chi sites to the left, to promote DNA circularization.

Upon injection into the next (recipient) cell, the terminal repeats must recombine to form circular DNA (Sternberg and Hoess, 1983). This DNA can either undergo rolling circle replication in a lytic infection or be replicated as a circular plasmid in a lysogenic infection, the two lifestyles of P1. Circularization requires the host RecBCD pathway: P1 replication and phage yields are reduced ∼20-fold and the phage do not form visible plaques in *recA* or *recBC* null mutants or in *recC* mutants altered in the RecBCD tunnel in which Chi is recognized (Zabrovitz et al., 1977; Amundsen et al., 2016). Thus, Chi-stimulated recombination is essential for P1 growth, likely at the circularization step. The abundant Chi sites, distributed as they are, ensure that RecBCD entering from either end of the injected linear DNA encounters an actively oriented Chi site. [Chi is active only when RecBCD approaches 5’ GCTGGTGG 3’ from the right as written here (Faulds et al., 1979; Taylor et al., 1985).] It is not surprising, then, that Chi is the most frequent octamer among the 44458 (top strand) or 55659 (both strands) octamers present in P1.

Several observations lead us to propose that Chi’s high frequency and distribution in the genome of P1 and its related phages reflect Chi being selected during evolution as a hotspot to promote recombination (circularization of the linear DNA after injection; Figure 7). This differs from the situation in *E. coli*, where we infer that selection for codon usage results in Chi’s high frequency. The following observations support this proposal.

1. P1 requires RecA-RecBCD-promoted recombination for growth, but *E. coli* does not.
2. Chi is located preferentially near the ends of P1’s genome, where recombination is required, but Chi is nearly uniformly distributed across *E. coli*’s genome, like its ORFs.
3. Chi is the most frequent octamer in P1 but is only the 21^st^-most frequent in *E. coli* (Table 1), suggesting a stronger or more direct selection for Chi in P1 than in *E. coli*.
4. Chi is much more dense in non-coding DNA (non-ORFs) in P1 than in *E. coli* (p = 0.0041; Table S6). In P1 Chi occurs at nearly the same density in ORFs and non-ORF sequences (p = 0.44), but it is 13 times more frequent in *E. coli* ORFs than non-ORFs (p < 0.0001; Table S6).
5. When Chi is in ORFs, the coding strand contains 5’ GCTGGTGG 3’ less often in P1 (63%) than in *E. coli* (79%) (p = 0.015; Table S6), indicating a closer connection between Chi and translation in *E. coli* than in P1.
6. CTG, part of this sequence, is the most frequent codon in *E. coli* but is only the fourth most frequent codon in P1 (Tables 3 and S7). The frequency of CTG, encoding Leu, is 3.2 times the mean of all codon frequencies in *E. coli* but only 1.7 times the mean in P1 (p < 0.0001 by two-tailed chi-square test), even though Leu is nearly as frequent in P1 (8.7%) as in *E. coli* (10.7%); Leu is the most frequent amino acid in *E. coli* and the second most frequent in P1 (Tables 3, S3, and S7). CTG is 50% of all Leu codons in *E. coli* but only 32% in P1 (p < 0.0001 by two-tailed chi-square test). Other codons are used at more nearly equal frequencies in the two genomes (Figure S7).
7. In *E. coli*, 68% of all Chi sites in ORFs are in the reading frame 5’ CTG|GTG 3’, significantly more than in P1 (41%, p < 0.0003 by two-tailed Fisher’s exact test; Tables 5 and S8), another indication that Chi is more closely associated with coding in *E. coli* than in P1.

Phages P7 (a close P1 relative) and D6 and putative phage NGUA22 [each distantly related to P1 and to each other (Gilcrease and Casjens, 2018)], also share these features with P1 – Chi is the most frequent octamer in their genomes, it is more abundant near their genome ends, it is nearly randomly oriented on their genomes, they likely require RecA and RecBCD to circularize their genomes for growth, and they use codon CTG about half as frequently as *E. coli* does. In other words, P1 and its relatives put Chi toward the genome ends (for recombination), and *E. coli* puts it in ORFs (for protein coding). This proposal and evidence for it are, to our knowledge, the first pointing toward a direct selection during evolution for Chi as a recombination hotspot.

### A myth: “Chi converts RecBCD from phage destruction to DNA repair”

This statement (Cheng et al., 2020) and similar ones so often written would be compatible with the dogma discussed above if it were true. But the mere growth of phage without Chi (Table S1) shows that phages without Chi are not destroyed by RecBCD. Among known phages, this view could not hold, for to our knowledge only *recBCD^+^* bacteria have been used as hosts to search for *E. coli* phages (G. Hatfull, pers. comm.). If RecBCD destroyed a phage, it would not have been observed and described. (It might be interesting, however, to search for phages that do grow only in *recBCD* mutants. Their behavior would almost certainly lead to new insights into phage and RecBCD physiology.)

This dogma likely stems from purified RecBCD’s activity at low ATP concentrations (less than that of Mg^2+^ ions). ATP chelates Mg^2+^; unchelated (“free”) Mg^2+^ is required for high RecBCD nuclease activity (Rosamond et al., 1979). With excess ATP, the free Mg^2+^ concentration is low and the nuclease activity is low; with excess Mg^2+^, the nuclease activity is high (Wright et al., 1971; Murphy, 2000). Thus, with excess ATP, RecBCD unwinds DNA from an end and simply nicks DNA at Chi, about five nucleotides to the 3’ side of 5’ GCTGGTGG 3’ (Taylor et al., 1985; Taylor and Smith, 1995) (Figure 1). With excess Mg^2+^, RecBCD frequently nicks DNA before encountering Chi [forming single-stranded (ss) DNA a few kb long], nicks at and near Chi (within and to both sides of 5’ GCTGGTGG 3’), nicks the complementary strand within and to both sides of 5’ CCACCAGC 3’, and nicks that strand during continued unwinding (Dixon and Kowalczykowski, 1993; Taylor and Smith, 1995; Anderson and Kowalczykowski, 1997a). Deducing which reaction occurs in *E. coli* cells is limited by lack of knowledge of the effective (free) concentration of Mg^2+^ in cells. Although in cells total Mg^2+^ (∼100 mM) is greater than total ATP (∼3 mM), most of the Mg^2+^ is likely bound to ribosomal RNA [see (Taylor and Smith, 1995; Smith, 2012)]. The effective (free) concentration of Mg^2+^ in *E. coli* cells is uncertain.

Genetic evidence, coupled with the biochemical activities of purified RecBCD, strongly indicates that <u>in *E. coli* cells RecBCD nicks DNA at Chi</u>, as purified RecBCD does with excess ATP. The activity of Chi is strongly influenced by nucleotides 4, 5, and 6 to the 3’ side of the Chi octamer 5’ GCTGGTGG 3’ (Amundsen et al., 2016; Taylor et al., 2016). This is true for genetic experiments in cells; it is also true for biochemical experiments with purified RecBCD when ATP is in excess but <u>not</u> when Mg^2+^ is in excess. Additional evidence also indicates simple nicking at Chi in cells (Smith, 2012). For example, Chi acts in *trans* (*i.e.,* between DNA molecules) in cells (Köppen et al., 1995; Myers et al., 1995) and with purified components with excess ATP but <u>not</u> with excess Mg^2+^ (Taylor and Smith, 1992, 1999). Furthermore, in lambda single-burst analyses, RecBCD pathway recombination is reciprocal (Sarthy and Meselson, 1976): ++ crossed with *ab* produces both +*b* <u>and</u> *a*+ at nearly equal frequencies in one infected cell. When Chi is between *a* and *b* and stimulates their recombination, degradation up to Chi, as envisaged by the dogma, would destroy one of the flanking alleles and not allow formation of reciprocal recombinants. Others report Chi-promoted recombination to be either reciprocal (Stahl et al., 1982; Kobayashi et al., 1984; Stahl et al., 1984), non-reciprocal (Lam et al., 1974; Stahl et al., 1980), or both (Ennis et al., 1987; Stahl et al., 1990; Stahl et al., 1995). Both reciprocal and non-reciprocal recombination promoted by RecBCD and Chi can be accounted for by the nick-at-Chi model (Figure 1). We are unaware of corresponding, consistent arguments for degradation up to Chi <u>in living cells</u>. Thus, the available evidence strongly supports simple nicking at Chi.

### An observation: RecBCD destroys DNA in *E. coli* only if it cannot recombine

The dogma holds that RecBCD destroys DNA that does not contain Chi. The growth of phages without Chi or RecBCD inhibitors, discussed above, indicates this view is incorrect. Additional evidence comes from Chi’s stimulation of HMW plasmid DNA in *E. coli*, noted above (Dabert et al., 1992). Formation of HMW DNA requires RecA DNA strand-exchange protein, an essential recombination factor that acts after RecBCD and Chi (Figure 1). RecA binds ss DNA and promotes formation of joint molecules with homologous ds DNA; joint molecules are resolved into recombinants, the products of DNA break repair. Purified RecBCD loads RecA onto the ss DNA produced by RecBCD after it cuts at Chi (Anderson and Kowalczykowski, 1997b). When RecBCD encounters properly oriented Chi on the “tail” of a rolling circle, the RecA-ssDNA complex can form a joint molecule with another plasmid in the cell, which can lead to a dumbbell-shaped plasmid without ends (Smith, 2012). The plasmid is thus protected from cellular exonucleases, including the dozens of other RecBCD molecules in the cell. Continued replication produces HMW DNA. Without RecA, RecBCD can still nick DNA at Chi on the rolling circle, but without RecA’s formation of joint molecules and recombinants, other RecBCD molecules or nucleases eventually destroy the DNA. [On ss DNA, RecBCD nuclease is highly active (Wright et al., 1971) but does not respond to Chi (Ponticelli et al., 1985).] Thus, Chi’s stimulation of HMW DNA formation cannot be taken as evidence that Chi switches RecBCD from a destructive nuclease to a beneficial repair enzyme. Coupled with RecA, the statement is true, but it is RecA-dependent recombination, not just Chi recognition by RecBCD, that blocks DNA destruction.

A different assay of DNA in *E. coli* leads to the same conclusion. Survival of plasmid DNA cut at a unique site is increased by a properly oriented Chi site and even more by two, tandem Chi sites (Kuzminov et al., 1994). Survival depends on RecA and ss DNA binding protein (SSB), which aids RecA in formation of joint DNA molecules. The surviving DNA contains branched DNA, likely arising from joint DNA molecule formation by RecA and SSB. If Chi inactivated RecBCD nuclease, this requirement for RecA would not be expected. As noted above, it is recombination, not just Chi, that prevents DNA destruction. In addition, plasmids form concatemers in recombination-proficient *recD* mutants, which lack RecBCD nuclease activity (Amundsen et al., 1986), only if RecA is present (Biek and Cohen, 1986), implying that other nucleases also can destroy DNA that does not recombine.

Growth of a phage is often assayed by plaque formation, which arises from a single phage particle infecting a cell. In this case, there is no homologous DNA with which the phage, or linearized form of it, can recombine to prevent eventual destruction. For example, RecBCD rapidly solubilizes about 40% of lambda DNA at a multiplicity of infection (MOI) of 1, a condition under which about 40% of the cells have no homolog with which lambda can recombine (Simmon and Lederberg, 1972). *I.e.*, a second (intact) DNA and presumably recombination are required for protection. Furthermore, degradation by RecBCD is at least as rapid on DNA with Chi as on DNA without Chi. Thus, clear evidence for Chi converting RecBCD from a destructive mode to a repair mode in cells is lacking.

### Mechanisms employing RecBCD that do distinguish self *vs.* non-self DNA

If Chi does not enable bacteria to tell self from non-self DNA, are there mechanisms that do? Yes – several mechanisms have been described, including some involving RecBCD but not in the manner envisaged by the dogma. [See (Rostol and Marraffini, 2019) for additional mechanisms not involving RecBCD.]

The first-described mechanism of self vs. non-self discrimination is restriction and modification by enzymes now universally used for DNA manipulation (Murray, 2002). In initial observations, phage T2 grown in *E. coli* strain X produced only a few phage that grew in another strain Y (*i.e.,* it was restricted by strain Y), but the rare phage that had grown in strain Y grew in both strain X and strain Y (*i.e.,* it was modified by strain Y); similar observations were made with other phages and bacterial strains (Luria, 1953). We now know that nearly all bacteria modify their own DNA and that of lucky phage escapees by methylation of the DNA at special sequences. Restriction enzymes cut DNA that is not modified (not methylated, non-self DNA) at these sites but do not cut modified (methylated, self) DNA. As noted above, phage lambda cut by *Eco*KI restriction enzyme is further degraded by RecBCD, whether or not it contains Chi (Simmon and Lederberg, 1972).

A more complex mechanism, CRISPR-Cas, was described more recently (Hille et al., 2018). About half of bacterial genomes examined contain Clustered, Regularly Interspersed, Palindromic Repeats (CRISPR). The repeats are ∼30 bp long and can form a “snap-back” internal hairpin; they are separated by ∼35 bp of unique sequence DNA (spacers). These “islands” usually contain <50, but occasionally >500, such alternating repeats and spacers. They remained mysterious for about 20 years after their discovery in *E. coli* [see (Ishino et al., 1987; Ishino et al., 2018)], when it was noted that the spacer DNA sequences were often that of phages or plasmids (*i.e.*, foreign or non-self DNA) (Bolotin et al., 2005; Mojica et al., 2005; Pourcel et al., 2005). This observation gave rise to the idea that CRISPR islands are involved in phage defense, or self vs. non-self recognition. Further experiments in *Streptococcus thermophilus* showed directly that infecting phage DNA could be integrated as new spacers in the CRISPR locus and that plasmid or phage DNA could be cut at a predetermined site upon transformation or infection (Barrangou et al., 2007; Garneau et al., 2010). Part of an RNA transcript of the CRISPR locus binds to a DNase, such as Cas9, and directs it to cleave the DNA at the position homologous to the RNA, itself homologous to the inserted (“foreign”) DNA. These observations led to new methods for genetic engineering, such as the insertion or deletion of DNA or change of a gene’s nucleotide sequence, at virtually any predetermined location by providing an appropriate RNA to activate the cleavage enzyme, usually Cas9 (Doudna and Charpentier, 2014; Lander, 2016).

Acquisition of new DNA spacers into the CRISPR locus requires the Cas1-Cas2 protein complex, which binds ds DNA ∼33 nucleotides long and can insert it into the CRISPR locus (Nunez et al., 2015). In cells, an additional repeat is generated concurrently. The mechanism by which this ds DNA substrate is generated is not clear, but in *E. coli* it (or another step leading to insertion) is regulated by RecBCD and Chi. During a 16-hour induction of Cas1-Cas2, *E. coli* (*i.e.*, self) DNA spacers are inserted into the CRISPR locus (Levy et al., 2015). (The strain used lacked Cas9, so cells with self DNA at the CRISPR locus survived.) Deep sequencing shows that spacer acquisition is reduced by ∼30% to the 5’ side of Chi (5’ GCTGGTGG 3’), the side to which Chi stimulates recombination (Figure 1); as expected, no Chi effect is observed in *recBCD* deletion mutants. These data indicate that Chi reduces acquisition of self DNA. Curiously, however, the frequency of acquisition is reduced by ∼50 – 70% in *recBCD* deletion mutants, indicating that RecBCD is actively involved in acquisition. In addition, acquisition to the 3’ side of Chi, and between Chi and an introduced ds DNA break, indicates that DNA encountered by RecBCD before Chi is not destroyed but, rather, is actively inserted into the CRISPR locus. One possibility is that ss DNA not bound by RecA is an important precursor to CRISPR insertions. If so, acquisition to the 3’ side of Chi, where RecBCD makes ss DNA, but not to the 5’ side of Chi, where RecBCD loads RecA, is accounted for.

Notably, the results of Levy et al. (Levy et al., 2015) show that self DNA is acquired; *i.e., E. coli* DNA is inserted into the *E. coli* CRISPR locus, at least in the absence of destructive Cas proteins. Presumably, these bacteria commit suicide when, as in wild-type cells, they contain the destructive Cas proteins, which would be directed to cleave the *E. coli* genome locus from which the newly acquired DNA came. These bacteria would be removed from the population, leaving only bacteria in which non-self DNA had been acquired. This scenario seems feasible, because the frequency of acquisition appears to be quite low – perhaps one bacterium in 10,000 inserts new DNA into the CRISPR locus per cell generation. If phage DNA were inserted into the CRISPR locus, these cells, though rare, would survive subsequent infection by that phage, whereas those that did not insert that phage DNA into the CRISPR locus would be killed by the phage. Eventually, only bacteria with the “correct” insert into the CRISPR locus would survive, and the population would survive subsequent phage infection. Thus, the apparent preference for non-self DNA being in the CRISPR locus could reflect random insertion followed by self destruction, a scenario opposite to that usually considered for CRISPR.

Another type of DNA island includes retrons, which encode a bacterial reverse transcriptase (RT) (Simon et al., 2019). One recently described retron, Ec48, engages RecBCD for its action – suicide of a phage-infected cell before production of mature phage to spare the rest of the bacterial population from infection and death (Millman et al., 2020). The Ec48 retron-encoded RT makes a 48-nucleotide DNA copy of part of the mRNA synthesized from the retron locus (Mao et al., 1997). The 2’ OH of a highly conserved G residue in the RNA primes the DNA synthesis, resulting in covalent linkage of DNA and RNA. Host RNase H removes most of the RNA, leaving a DNA-RNA complex of ∼170 nucleotides. When a phage that encodes a RecBCD inhibitor, such as lambda Gam, infects the cell, the inhibited RecBCD and the DNA-RNA complex activate an “effector” protein, also encoded by the retron, to kill the cell (Millman et al., 2020). The Ec48 effector protein contains two putative transmembrane regions, suggesting that activation of this protein makes the cell membrane leaky and leads to death. Although Chi has no known role in Ec48 action, RecBCD is clearly involved, in an unknown manner, in telling the cell that non-self DNA has entered the cell and it’s time to commit suicide and thus protect the population.

### Conclusion: Telling self from non-self is complex

Given that the basic genetic material in all cells and plasmids and many phages is the same – DNA – it is no surprise that cells have a hard time telling self from non-self on the basis of DNA. Even in cases where bacteria have evolved this discrimination, many phages have won the “arms race” by evolving anti-restriction and anti-CRISPR mechanisms (Tock and Dryden, 2005; Stanley and Maxwell, 2018). From this viewpoint, it is understandable that something as simple as Chi does not do the trick. Nevertheless, the stimulation of a critical function – DNA break repair and genetic recombination – by Chi appears widely important for cells, because the Chi-RecBCD combination appears universal in enteric bacteria, and related cohorts may exist in many other bacterial species. The intersection of RecBCD with recombination, CRISPR spacer acquisition, and retrons may reflect RecBCD’s central role in maintaining life.

### Methods of data analysis

Reference numbers for genome sequences are in Table S9. Scripting for computational analysis of genomic data was done chiefly in Python 3.9. Statistical tests and linear regression analyses used Graphpad and Excel.

## Acknowledgements

We are grateful to Sue Amundsen, Graham Hatfull, Eduardo Rocha, Lyle Simmons, Rasi Subramaniam, and especially Sherwood Casjens for helpful discussions; and Sue Amundsen, Justin Courcelle, Gareth Cromie, Kevin Forsberg, Khoi Ha, and Yihua Zhu for comments on the manuscript. This research was supported by MIRA grant R35 GM118120 from the National Institute of General Medical Sciences of the United States of America to GRS.

## Supplemental Information

**Figure S1.**
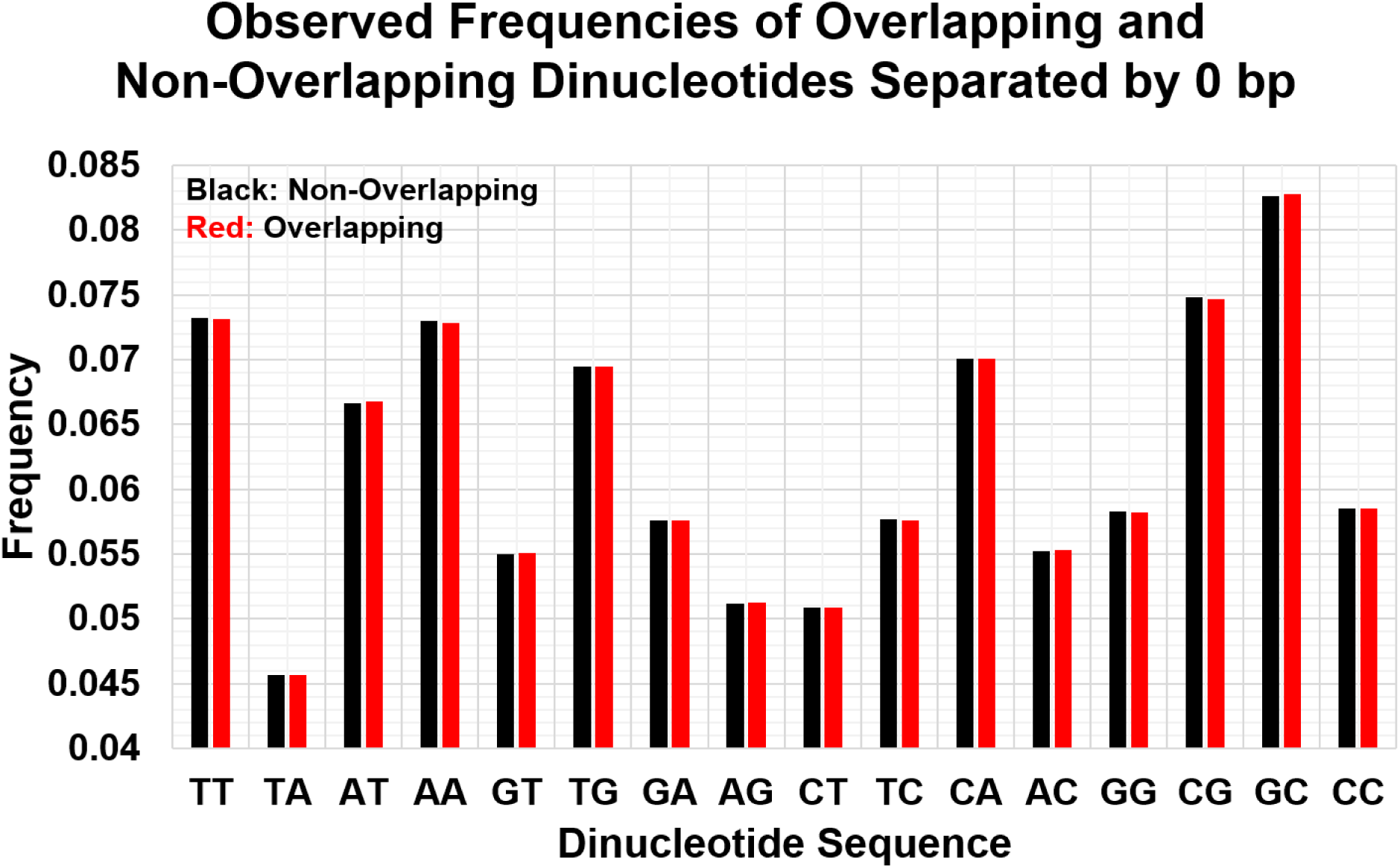
Overlapping and non-overlapping dinucleotide frequency patterns are indistinguishable. Observed frequencies of each dinucleotide on the top *E. coli* strand were enumerated by counting adjacent nucleotides without overlap (1 2, 3 4, 5 6, …; black) or with overlap (1 2, 2 3, 3 4, …; red). The patterns are not significantly different (p > 0.999 by unpaired *t* test). Red data are also in Figure 1A.

**Figure S2.**
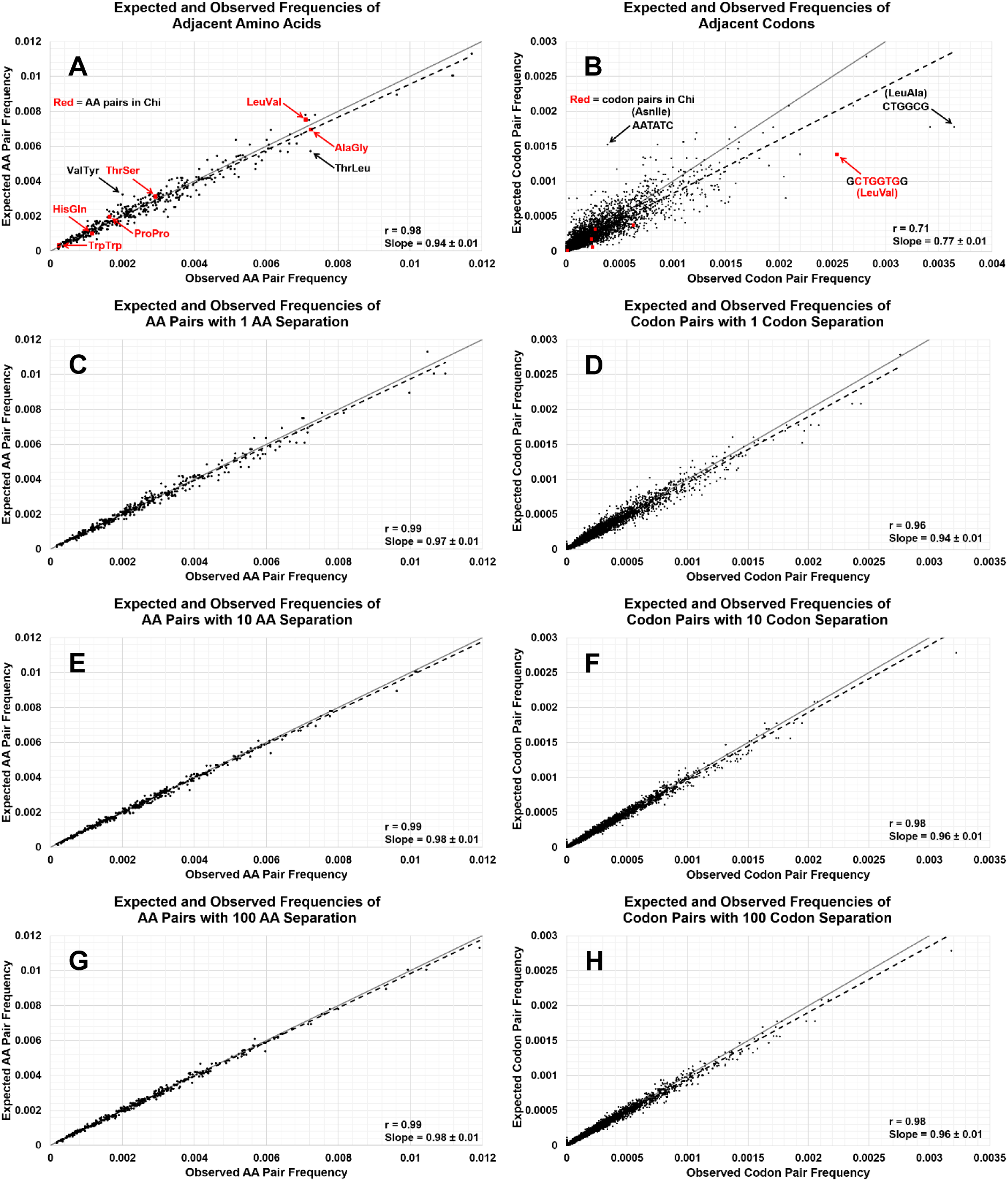
Adjacent *E. coli* codon pairs, like amino acid pairs, are not randomly distributed, but their distributions become progressively more nearly random when separated by 1, 10, or 100 codons (or amino acids). (**A**) The observed frequency of each dipeptide is plotted against its expected frequency (product of individual amino acid frequencies in Table S3). Note that the slope of the linear regression line (dashed black) is significantly different from 1 (p < 0.0001 by two-sided unpaired t test), which would indicate randomness. In red are dipeptides encoded within Chi or its complement. In black are two dipeptides whose observed frequencies are especially different from their expected frequencies. Data are also in Figure 4. (**B**) The corresponding analysis for dicodons, from Figure 5. In red are codons within Chi or its complement. In black are two dicodons whose observed frequencies are especially different from their expected frequencies. (**C – H**) Corresponding analyses for dipeptides and dicodons separated by 1 (**C** and **D**), 10 (**E** and **F**), or 100 (**G** and **H**) amino acids or codons. Note that the correlation coefficients **r** and slopes approach 1, indicating randomness, as the separation increases from 0 to 1 to 10 but not further with separation of 100.

**Figure S3.**
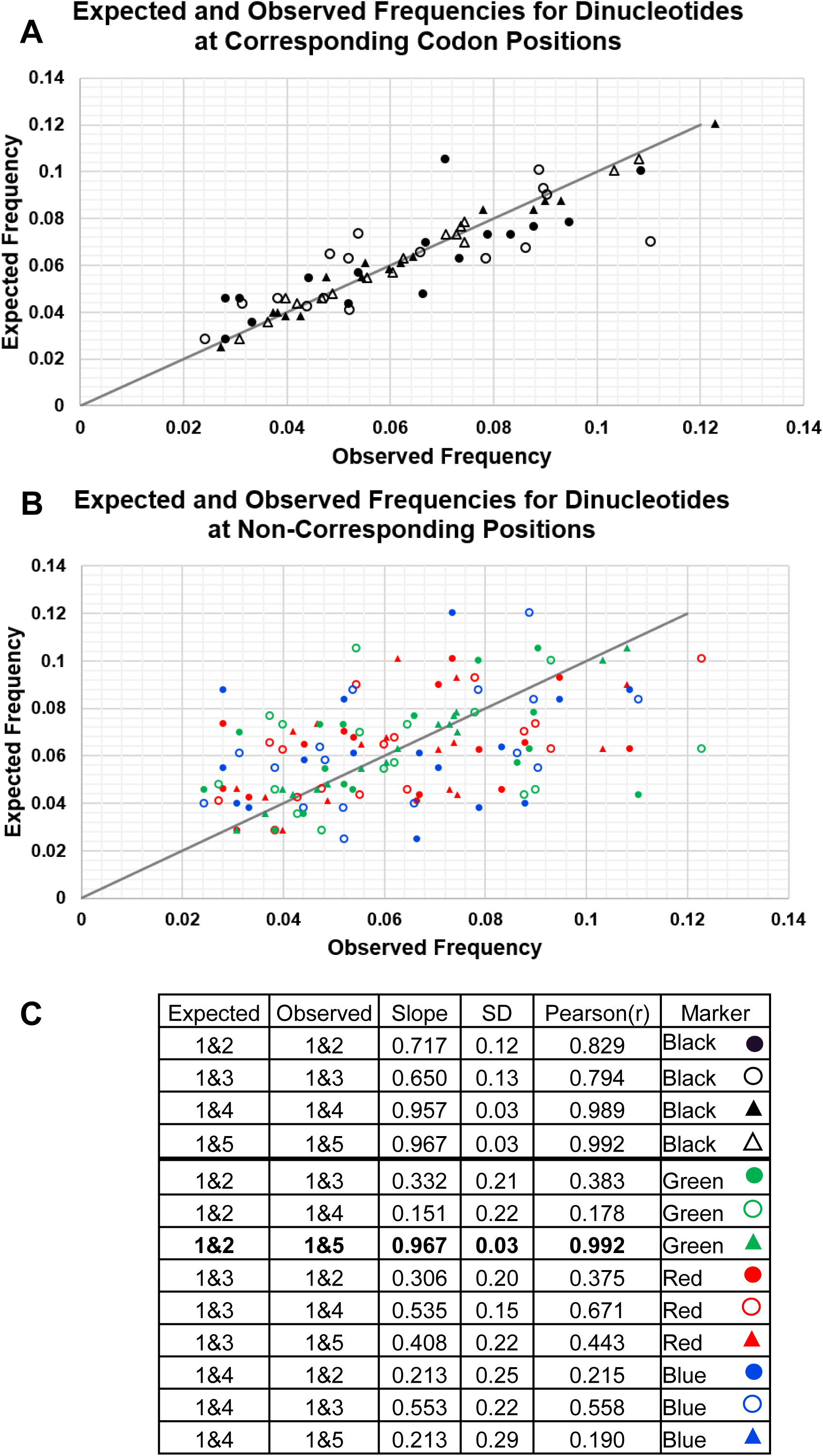
Nucleotides separated by 0 to 3 nucleotides follow patterns set by *E. coli*’s codon usage. (**A** and **B**) The frequencies of nucleotides at each position in codons (Table 3) were used to calculate expected frequencies of nucleotide pairs in the positions in codons (indicated in panel **C**) within each ORF and plotted *vs.* the observed frequencies. Lines indicate equality. (**C**) Data for the same separation distance (“corresponding”; top four lines and graphed in panel **A**) are more highly correlated (slope of linear regression and Pearson correlation coefficient **r** are closer to unity) than data for different separation distances (“non-corresponding”; bottom nine lines and graphed in panel **B**). An exception is 1&2 compared with 1&5 (bold type), which compares the first and second nucleotides in a codon with the first nucleotide in a codon and the second nucleotide in the adjacent codon; as noted in Figure 2, these are correlated.

**Figure S4.**
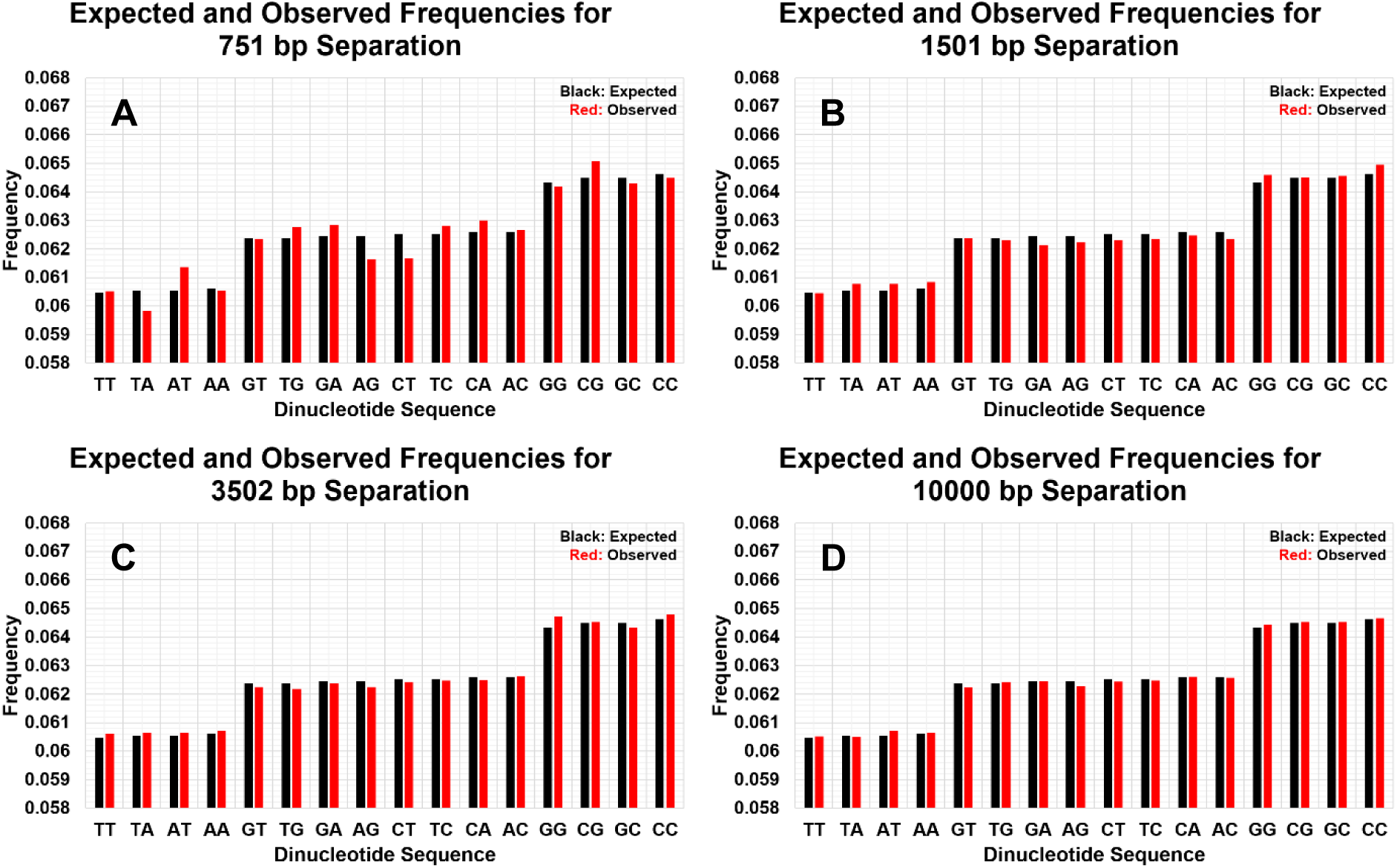
Frequencies of *E. coli*’s nucleotide pairs approach those expected from random association of nucleotides as the distance between the nucleotides in the pair approaches and exceeds the length of ORFs. Expected frequencies of each nucleotide pair based on random association of nucleotides and their frequencies on the top strand of *E. coli* (Table S2) are shown in black. Observed frequencies of nucleotide pairs separated by 751 (**A**), 1501 (**B**), 3502 (**C**), or 10000 (**D**) nucleotides on the top strand of *E. coli* are shown in red. These separations place the nucleotides in the same positions (denoted 1&3, 2&1, and 3&2 in text related to Figure 1C - F) of codons in the same ORF or about 1/3 of the time if they are in different ORFs. See Figure 2 for ORF distribution lengths. Panel D is also in Figure 2B.

**Figure S5.**
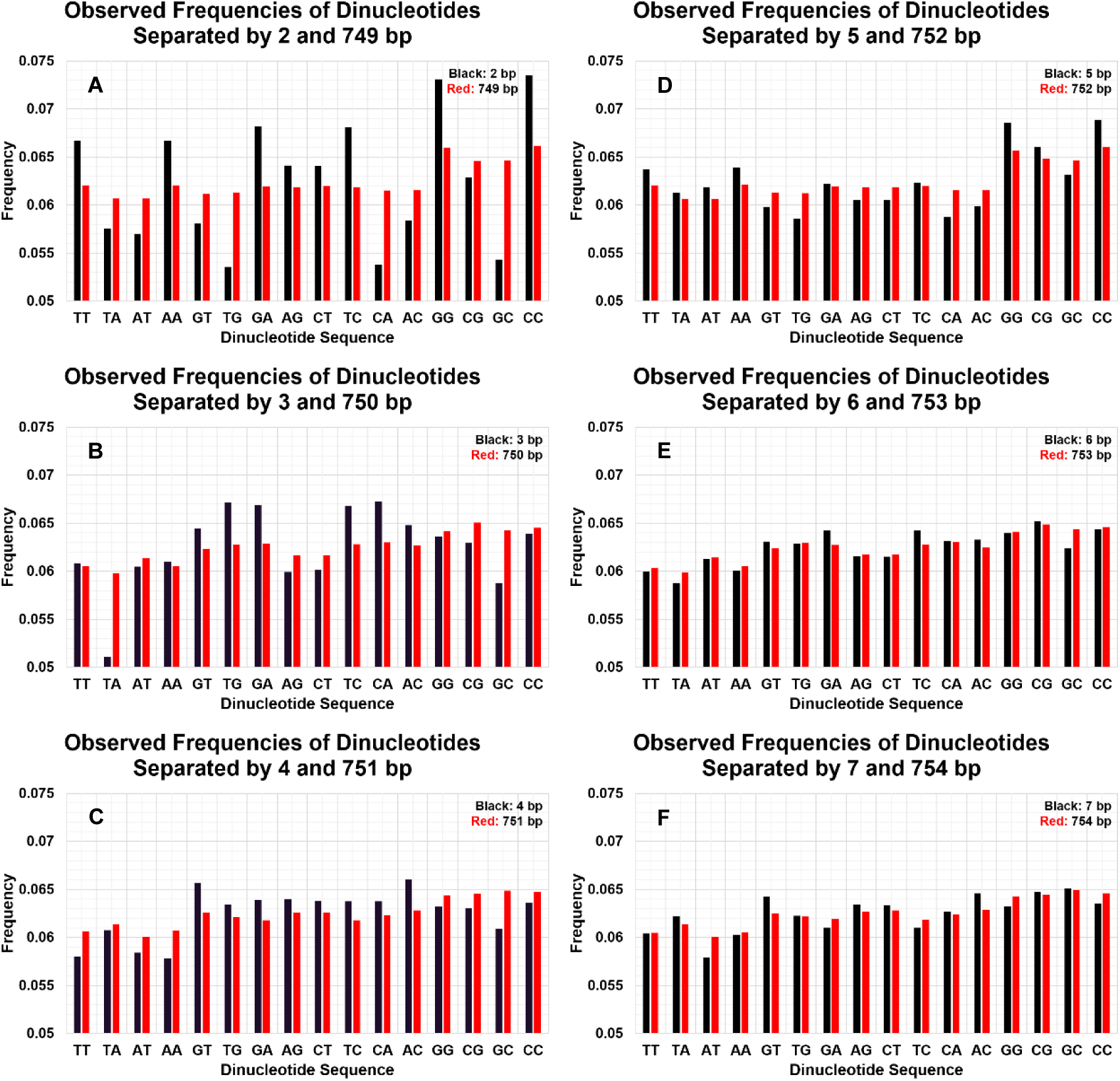
Nucleotides separated by even 750 bp are non-random and display a 3-nucleotide repeating pattern. The top two panels show frequencies of nucleotide pairs on *E. coli*’s top strand separated by 2 and 749 (**A**) or 5 and 752 (**D**) nucleotides, which places them at the same positions in codons if the nucleotide pair occurs within an ORF. A similar relation holds for panels B and E, with nucleotides one position displaced in codons, and for panels C and F, with nucleotides two positions displaced in codons. The pattern of nucleotides separated by 749 nucleotides (**A**) is nearly the same as that separated by 752 nucleotides (**D**) but differs from those separated by 750 (**B**) or 751 (**C**) nucleotides. The frequencies approach that of random association (Figures 2B and S4D) as the distance between the nucleotides in the pair increases.

**Figure S6.**
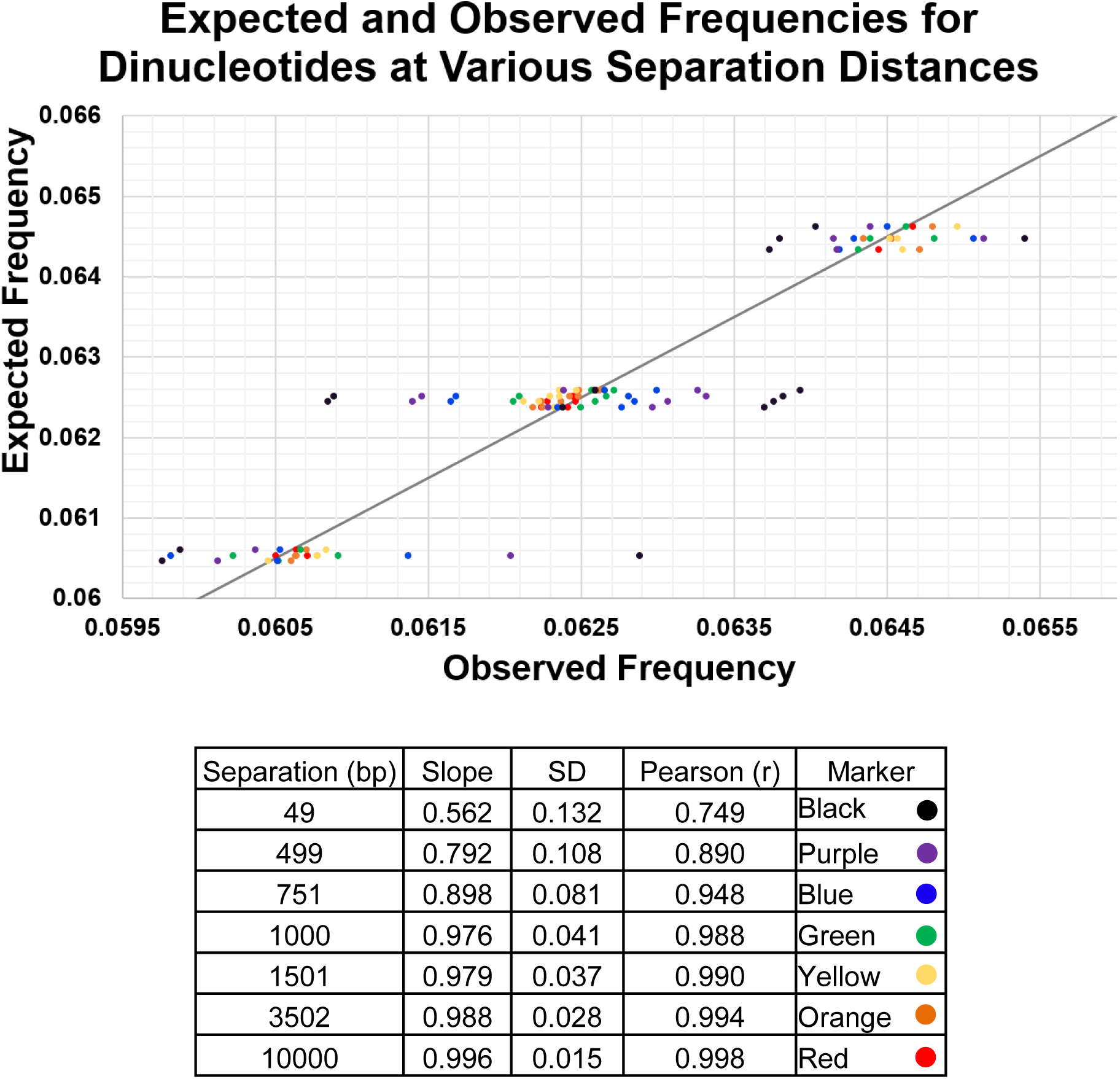
Frequencies of nucleotide pairs (top strand) approach those expected from random association of nucleotides as the distance between the nucleotides in the pair approaches and exceeds the length of ORFs (Figure 2). Linear regression analysis used the observed frequencies of nucleotide pairs separated by the indicated distances on the top *E. coli* strand and the expected frequencies from random association of nucleotides (Table S2). Slopes ± SD and Pearson’s correlation coefficient **r** were tabulated. See Figures 2A, 2B, S4, and S5 for corresponding data with separations of 0 – 7, 749 – 754, 1501, 3502, and 10000 bp and Figure 3 for ORF distribution lengths.

**Figure S7.**
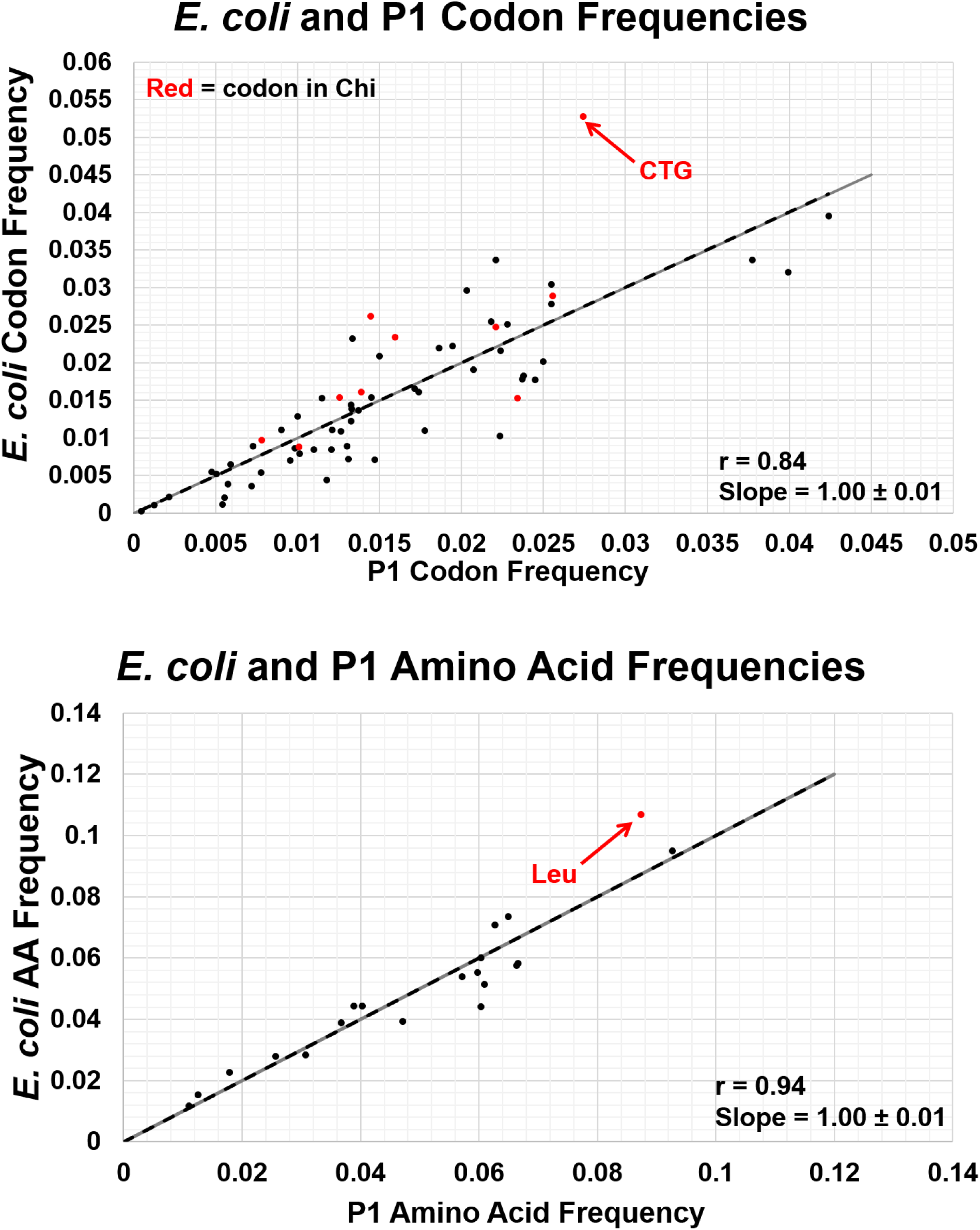
Leu and its codon CTG are less frequent in P1 than in *E. coli*. The frequencies of codons (top) and encoded amino acids (bottom) in *E. coli* are plotted *vs.* their frequencies in phage P1. Note that the slope of the linear regression line (dashed black) is indistinguishable from 1 (solid grey line), indicating equal frequencies in the two genomes. Pearson correlation coefficient **r** is near unity, indicating high correlation. But in red are two outliers – Leu and its most frequent codon, CTG, which occurs in Chi.

**Table S1.** Chi is present in many *E. coli* phages and conjugative plasmids.

| *E. coli* phage or<br>plasmid | # Chi<br>sites | Genome<br>Length (bp) | # Chi/Mb | % G + C |
| --- | --- | --- | --- | --- |
| Lambda and 17 related<br>phages* | 0 | (~)48502 | 0 | 49.9 |
| Phage P22<br>(*Salmonella*, λ-like)* | 0 | 41733 | 0 | 47.1 |
| HK639 (lambda-like)** | 13 | 49576 | 262 | 52.0 |
| phi80 (lambda-like)** | 9 | 46150 | 195 | 52.1 |
| 2H10 (lambda-like)*** | 8 | 46685 | 171 | 50.3 |
| HK225 (lambda-like)*** | 7 | 45366 | 154 | 52.0 |
| T3 (virulent) | 12 | 38208 | 314 | 49.9 |
| T5 (virulent) | 8 | 121750 | 65 | 39.3 |
| T7 (virulent)** | 7 | 39937 | 175 | 48.4 |
| P1 (temperate) | 50 | 94800 | 527 | 47.3 |
| P7 (P1-like) | 51 | 101660 | 501 | 47.4 |
| D6 (P1-like) | 53 | 91159 | 581 | 47.0 |
| NGUA22 (P1-like)**** | 68 | 93583 | 727 | 49.4 |
| F factor plasmid | 0 | 99159 | 0 | 48.2 |
| Plasmid pJP4 | 36 | 87688 | 411 | 64.7 |
| Plasmid pMG252 | 81 | 185643 | 436 | 52.8 |
| Plasmid pR100 | 3 | 94281 | 31 | 52.0 |
| Plasmid pR6K | 1 | 39872 | 25 | 45.3 |

\* Has RecBCD-inhibitor (Gam, Abc2) (Sakaki et al., 1973; Poteete et al., 1988)
\*\* Has putative RecBCD-inhibitor (GamL, gp5.9) (Qimron et al. 2010; Pacumbaba and Center, 1975; Rotman et al., 2012)
\*\*\* Has no obvious RecBCD-inhibitor (Bobay et al., 2013).
\*\*\*\* NGUA22 is a *Salmonella enterica* plasmid with features like P1 and D6 prophages suggesting that it is the prophage form of a phage (Gilcrease and Casjens, 2018).
Data are from DNA sequences whose sources are in Table S9.

**Table S2.** Top strand nucleotide frequencies in *E. coli* and *S. aureus*.

| Nucleotide | *Escherichia coli* |  | *Staphylococcus aureus* |  |
| --- | --- | --- | --- | --- |
|  | Number | Frequency | Number | Frequency |
| Thymine (T) | 1141382 | 0.2459 | 955315 | 0.3386 |
| Adenine (A) | 1142742 | 0.2462 | 938713 | 0.3327 |
| Guanine (G) | 1177437 | 0.2537 | 461500 | 0.1636 |
| Cytosine (C) | 1180091 | 0.2542 | 465832 | 0.1651 |
| Total | 4641652 | 1 | 2821360 | 1 |

Data are from DNA sequences whose sources are in Table S9.

**Table S3.** The four most abundant.

| Amino acid* | Number | Frequency | Amino acid | Number | Frequency |
| --- | --- | --- | --- | --- | --- |
| **(L) Leucine** | 145018 | 0.1067 | **(A) Alanine** | 2607 | 0.0927 |
| **(A) Alanine** | 129054 | 0.0949 | **(L) Leucine** | 2457 | 0.0874 |
| **(G) Glycine** | 100067 | 0.0736 | (S) Serine | 1873 | 0.0666 |
| **(V) Valine** | 96345 | 0.0709 | (E) Glutamic Acid | 1867 | 0.0664 |
| (I) Isoleucine | 81749 | 0.0601 | **(G) Glycine** | 1827 | 0.0650 |
| (S) Serine | 79125 | 0.0582 | **(V) Valine** | 1762 | 0.0626 |
| (E) Glutamic Acid | 78143 | 0.0575 | (D) Aspartic Acid | 1714 | 0.0609 |
| (R) Arginine | 75250 | 0.0554 | (I) Isoleucine | 1697 | 0.0603 |
| (T) Threonine | 73400 | 0.0540 | (K) Lysine | 1697 | 0.0603 |
| (D) Aspartic Acid | 69827 | 0.0514 | (R) Arginine | 1681 | 0.0598 |
| (Q) Glutamine | 60318 | 0.0444 | (T) Threonine | 1608 | 0.0572 |
| (P) Proline | 60262 | 0.0443 | (N) Asparagine | 1324 | 0.0471 |
| (K) Lysine | 59939 | 0.0441 | (Q) Glutamine | 1133 | 0.0403 |
| (N) Asparagine | 53607 | 0.0394 | (P) Proline | 1092 | 0.0388 |
| (F) Phenylalanine | 52869 | 0.0389 | (F) Phenylalanine | 1033 | 0.0367 |
| (Y) Tyrosine | 38683 | 0.0285 | (Y) Tyrosine | 865 | 0.0308 |
| (M) Methionine | 37931 | 0.0279 | (M) Methionine | 720 | 0.0256 |
| (H) Histidine | 30842 | 0.0227 | (H) Histidine | 504 | 0.0179 |
| (W) Tryptophan | 20896 | 0.0154 | (W) Tryptophan | 355 | 0.0126 |
| (C) Cysteine | 15935 | 0.0117 | (C) Cysteine | 310 | 0.0110 |
|  | 1359260 | 1 |  | 28126 | 1 |

\* Letters in parentheses are the one-letter designation for the indicated amino acids.
Data are from Tables 3 and S7.

**Table S4.** “Chi” (5’ GAAGCGG 3’) contains *S. aureus*’s second most frequent codon GAA.

| Codon | Number | Codon | Number | Codon | Number | Codon | Number |
| --- | --- | --- | --- | --- | --- | --- | --- |
| †(F)TTT | 26378 | (S) TCT | 10390 | (Y) TAT | 24422 | (C) TGT | 4117 |
| (F) TTC | 9667 | (S) TCC | 1356 | (Y) TAC | 6829 | (C) TGC | 1035 |
| (L) TTA | 43151 | (S) TCA | 16217 | (*) TAA | 2071 | (*) TGA | 368 |
| (L) TTG | 10903 | (S) TCG | 3175 | (*) TAG | 453 | (W) TGG | 6040 |
| (L) CTT | 8628 | (P) CCT | 8571 | (H) CAT | 14721 | (R) CGT | 10536 |
| (L) CTC | 1607 | (P) CCC | 813 | (H) CAC | 3607 | (R) CGC | 2524 |
| (L) CTA | 6884 | (P) CCA | 12991 | (Q) CAA | 28853 | (R) CGA | 3929 |
| (L) CTG | 1923 | (P) CCG | 3232 | (Q) CAG | 4032 | (R) **CGG** | 383 |
| (I) ATT | 41950 | (T) ACT | 13394 | (N) AAT | 34454 | (S) AGT | 13359 |
| (I) ATC | 11856 | (T) ACC | 2145 | (N) AAC | 10887 | (S) **AGC** | 4160 |
| (I) ATA | 15171 | (T) ACA | 23082 | (K) AAA | 48783 | (R) AGA | 9360 |
| (M) ATG | 20785 | (T) ACG | 7601 | (K) **AAG** | 11370 | (R) AGG | 1190 |
| (V) GTT | 21813 | (A) GCT | 16137 | (D) GAT | 36304 | (G) GGT | 25859 |
| (V) GTC | 5886 | (A) GCC | 3537 | (D) GAC | 10029 | (G) GGC | 7406 |
| (V) GTA | 18111 | (A) GCA | 23839 | (E) **GAA** | 43396 | (G) GGA | 11261 |
| (V) GTG | 7614 | (A) **GCG** | 7482 | (E) GAG | 8403 | (G) GGG | 3490 |
| Total Codons: 799920 [2892 ORFs] |  |  |  |  |  |  |  |

† Letters in parentheses indicate the encoded amino acid, using the one-letter abbreviations in Table S3.
Data are for all ORFs ≥14 codons long as annotated in *S. aureus* strain NCTC 8325 (GenBank accession number NC\_007795.1). Bold-face codons are in “Chi.”

**Table S5.** Like E. coli Chi, S. aureus “Chi” frequency is correlated with its in-frame dicodon frequencies.

| Dicodon | Expected number* | Observed number | Proportion observed (%) | Proportion of "Chi" sites in this frame (%) |
| --- | --- | --- | --- | --- |
| GAAGCG | 409 | 268 | 82.5 | 82.6 |
| AAGCGG | 5 | 3 | 0.9 | 1.0 |
| Subtotal | 414 | 271 |  |  |
| CCGCTT | 35 | 25 | 7.7 | 3.3 |
| CGCTTC | 31 | 29 | 8.9 | 3.3 |
| Subtotal | 66 | 54 |  |  |
| Total | 480 | 325 | 100 | 90.2 |

\* Based on random codon association, calculated using values from Table S4.

**Table S6.** In *E. coli,* but not in P1, Chi is preferentially in ORFs.

| **Species:** | **P1** | **P7** | **D6** | **NGUA22** | ***E. coli*** |
| --- | --- | --- | --- | --- | --- |
|  | # (Fraction) | # (Fraction) | # (Fraction) | # (Fraction) | # (Fraction) |
| Top strand Chi<sup>a</sup> | 31/50 (0.62) | 32/51 (0.63) | 20/53 (0.38) | 24/68 (0.35) | 499/1008 (0.50) |
| Top strand Comp<sup>a</sup> | 19/50 (0.38) | 19/51 (0.37) | 33/53 (0.62) | 44/68 (0.65) | 509/1008 (0.50) |
| ORF Chi<sup>b</sup> | 29/46 (0.63) | 28/46 (0.61) | 38/52 (0.73) | 41/62 (0.66) | 792/997 (0.79) |
| **p\*** | **0.015** | **0.0051** | **0.29** | **0.0166** | **NA** |
| non-ORF Chi+Comp<sup>c</sup> | 4/50 (0.080) | 5/51 (0.098) | 1/53 (0.019) | 6/68 (0.088) | 11/1008 (0.011) |
| **p\*\*** | **0.0041** | **0.0006** | **0.461** | **0.0004** | **NA** |
| Non-ORF DNA | 10855/94800 (0.115) | 9831/101660 (0.0967) | 10057/91159 (0.110) | 9623/93583 (0.103) | 579575/4641652 (0.125) |
| **p\*\*\*** | **0.444** | **0.974** | **0.034** | **0.689** | **<0.0001** |

<sup>a</sup> Number of times Chi (5' GCTGGTGG 3') or its complement (Comp; 5' CCACCAGC 3') occurs on the top strand among all Chi sites on both strands.
<sup>b</sup> Number of times Chi (5' GCTGGTGG 3') occurs in ORFs among all occurrences of Chi or its complement in ORFs.
<sup>c</sup> Number of times Chi or its complement occurs in non-coding DNA among all Chi sites on both strands.
\* Probability (by two-tailed Fisher's exact test) that the fraction of Chi (5' GCTGGTGG 3') in ORFs is the same as that in *E. coli*.
\*\* Probability (by two-tailed Fisher's exact test) that the fraction of Chi plus its complement not in ORFs is the same as that in *E. coli*.
\*\*\* Probability (by two-tailed chi-square test) that the fraction of Chi plus its complement not in ORFs is the same as the fraction of DNA not in ORFs.

**Table S7.** P1 codons are less enriched for Chi sequences than are E. coli’s.

| Codon | Number | Codon | Number | Codon | Number | Codon | Number |
| --- | --- | --- | --- | --- | --- | --- | --- |
| (F)†TTT | 549 | (S) TCT | 341 | (Y) TAT | 491 | (C) TGT | 143 |
| (F) TTC | 484 | (S) TCC | 278 | (Y) TAC | 374 | (C) TGC | 167 |
| (L) TTA | 376 | (S) TCA | 370 | (*) TAA | 61 | (*) TGA | 36 |
| (L) TTG | 387 | (S) TCG | 206 | (*) TAG | 13 | (W) **TGG** | 355 |
| (L) CTT | 502 | (P) CCT | 270 | (H) CAT | 283 | (R) CGT | 423 |
| (L) CTC | 254 | (P) CCC | 135 | (H) CAC | 221 | (R) CGC | 526 |
| (L) CTA | 163 | (P) CCA | 310 | (Q) CAA | 410 | (R) CGA | 203 |
| (L) **CTG** | 775 | (P) CCG | 377 | (Q) CAG | 723 | (R) CGG | 219 |
| (I) ATT | 720 | (T) ACT | 368 | (N) AAT | 692 | (S) AGT | 285 |
| (I) ATC | 644 | (T) ACC | 451 | (N) AAC | 632 | (S) AGC | 393 |
| (I) ATA | 333 | (T) ACA | 415 | (K) AAA | 1066 | (R) AGA | 157 |
| (M) ATG | 720 | (T) ACG | 374 | (K) AAG | 631 | (R) AGG | 153 |
| (V) GTT | 672 | (A) **GCT** | 662 | (D) GAT | 1128 | (G) **GGT** | 625 |
| (V) GTC | 324 | (A) GCC | 616 | (D) GAC | 586 | (G) GGC | 574 |
| (V) GTA | 357 | (A) GCA | 705 | (E) GAA | 1197 | (G) GGA | 286 |
| (V) **GTG** | 409 | (A) GCG | 624 | (E) GAG | 670 | (G) GGG | 342 |
| Total Codons: 28236 [110 ORFs] |  |  |  |  |  |  |  |

† Letters in parentheses indicate the encoded amino acid, using the one-letter abbreviations in Table S3.
Data are for all ORFs $\geq 14$ codons long as annotated in phage P1 (GenBank accession number NC\_005856.1). Bold-face codons are in Chi (5' GCTGGTGG 3').

**Table S8.** Chi’s frequency is correlated with its in-frame dicodon frequencies in phage P1 and relatives less often than in *E. coli*.

**P1 Dicodon Analysis**
| Dicodon | Expected number* | Observed number | Proportion of observed (%) | Proportion of Chi sites in this frame (%) |
| --- | --- | --- | --- | --- |
| GCTGGT | 15 | 24 | 22.86 | 13.04 |
| CTGGTG | 12 | 29 | 27.62 | 41.30 |
| TGGTGG | 5 | 10 | 9.52 | 8.70 |
| Subtotal | 32 | 63 |  |  |
| CCACCA | 3 | 6 | 5.71 | 6.52 |
| CACCAG | 6 | 13 | 12.38 | 8.70 |
| ACCAGC | 6 | 23 | 21.90 | 21.74 |
| Subtotal | 15 | 42 |  |  |
| Total | 47 | 105 | 100 | 100 |

**P7 Dicodon Analysis**
| Dicodon | Expected number* | Observed number | Proportion of observed (%) | Proportion of Chi sites in this frame (%) |
| --- | --- | --- | --- | --- |
| GCTGGT | 17 | 33 | 27.50 | 13.04 |
| CTGGTG | 16 | 34 | 28.33 | 41.30 |
| TGGTGG | 5 | 9 | 7.50 | 6.52 |
| Subtotal | 38 | 76 |  |  |
| CCACCA | 4 | 8 | 6.67 | 8.70 |
| CACCAG | 5 | 13 | 10.83 | 6.52 |
| ACCAGC | 8 | 23 | 19.17 | 23.92 |
| Subtotal | 17 | 44 |  |  |
| Total | 55 | 120 | 100 | 100 |

**D6 Dicodon Analysis**
| Dicodon | Expected number* | Observed number | Proportion of observed (%) | Proportion of Chi sites in this frame (%) |
| --- | --- | --- | --- | --- |
| GCTGGT | 16 | 33 | 28.21 | 19.23 |
| CTGGTG | 12 | 37 | 31.62 | 48.08 |
| TGGTGG | 4 | 6 | 5.13 | 5.77 |
| Subtotal | 32 | 76 |  |  |
| CCACCA | 4 | 7 | 5.98 | 3.85 |
| CACCAG | 5 | 14 | 11.97 | 15.38 |
| ACCAGC | 5 | 20 | 17.09 | 7.69 |
| Subtotal | 14 | 41 |  |  |
| Total | 46 | 117 | 100 | 100 |

**NGUA22 Dicodon Analysis**
| Dicodon | Expected number* | Observed number | Proportion of observed (%) | Proportion of Chi sites in this frame (%) |
| --- | --- | --- | --- | --- |
| GCTGGT | 12 | 30 | 21.90 | 17.74 |
| CTGGTG | 16 | 51 | 37.22 | 43.55 |
| TGGTGG | 6 | 10 | 7.30 | 4.84 |
| Subtotal | 34 | 91 |  |  |
| CCACCA | 3 | 4 | 2.92 | 3.23 |
| CACCAG | 7 | 12 | 8.76 | 8.06 |
| ACCAGC | 8 | 30 | 21.90 | 22.58 |
| Subtotal | 18 | 46 |  |  |
| Total | 52 | 137 | 100 | 100 |

\* Based on random codon association, calculated using values from Table S7.

**Table S9.** Sources of nucleotide sequences.

| **Species** | **Identifier** | **Date Accessed** |
| --- | --- | --- |
| *Bacillus subtilis* | NC_000964.3 | 4/29/2021 |
| *Escherichia coli* | NC_000913.3 | 11/16/2020 |
| *Klebsiella pneumoniae* | NC_016845.1 | 12/14/2020 |
| *Mycobacterium tuberculosis* | NC_000962.3 | 2/23/2021 |
| *Pseudomonas aeruginosa* | NZ_LN831024.1 | 12/14/2020 |
| *Salmonella enterica* | NC_003197.2 | 2/23/2021 |
| *Serratia sp.* | NZ_CP032738.1 | 2/23/2021 |
| *Staphylococcus aureus* | NC_007795.1 | 12/14/2020 |
| *Vibrio harveyi* | NZ_CP009467.1 | 2/23/2021 |
| *Xanthomonas sp.* | NZ_LR861805.1 | 2/23/2021 |

| **Phage** | **Identifier** | **Date Accessed** |
| --- | --- | --- |
| Lambda | NC_001416.1 | 12/24/2020 |
| HK639 (lambda-like) | NC_016158.1 | 12/28/2020 |
| Φ80 (lambda-like) | JX871397 | 12/24/2020 |
| 2H10 (lambda-like) | LR595862 | 12/28/2020 |
| HK225 (lambda-like) | JQ086371 | 12/28/2020 |
| P1 (temperate) | NC_005856.1 | 11/16/2020 |
| P7 (P1-like) | NC_050152.1 | 1/21/2021 |
| D6 (P1-like) | MF356679.1 | 12/15/2020 |
| NGUA22 (P1-like) | BCOU01000029.1 | 6/20/2021 |
| T3 (virulent) | NC_003298.1 | 1/5/2021 |
| T5 (virulent) | NC_005859.1 | 1/16/2021 |
| T7 (virulent) | NC_001604.1 | 1/12/2021 |

| **Plasmid** | **Identifier** | **Date Accessed** |
| --- | --- | --- |
| F factor plasmid | AP001918.1 | 2/16/2021 |
| pJP4 | NZ_AY365053.1 | 2/16/2021 |
| pMG252 | NZ_MK638972.1 | 2/16/2021 |
| pR100 | NC_002134 | 2/16/2021 |
| pR6K | NZ_LT827129.1 | 2/16/2021 |

